# ILoReg enables high-resolution cell population identification from single-cell RNA-seq data

**DOI:** 10.1101/2020.01.20.912675

**Authors:** Johannes Smolander, Sini Junttila, Mikko S Venäläinen, Laura L Elo

**Affiliations:** Turku Bioscience Centre, University of Turku and Åbo Akademi University, Tykistökatu 6, 20520 Turku, Finland

## Abstract

Single-cell RNA-seq allows researchers to identify cell populations based on unsupervised clustering of the transcriptome. However, subpopulations can have only subtle transcriptomic differences and the high dimensionality of the data makes their identification challenging. We introduce ILoReg (https://github.com/elolab/iloreg), an R package implementing a new cell population identification method that achieves high differentiation resolution through a probabilistic feature extraction step that is applied before clustering and visualization.

## Main

Single-cell RNA-seq (scRNA-seq) enables identification of known and novel cell populations by unsupervised clustering of transcriptomic profiles of individual cells. However, the high number of genes presents a major challenge for the analysis of scRNA-seq data by increasing the similarity of distances between the cells, a phenomenon known as the ‘curse of dimensionality’^1^. To reduce its effect, scRNA-seq pipelines typically apply a feature selection step that selects a set of highly variable genes prior to unsupervised clustering^2^. However, this approach can eliminate genes that are important for the identification of the underlying cell populations of a sample or add unwanted variation if irrelevant features are chosen. Moreover, the number of remaining genes is typically still in the thousands and detecting cell populations with subtle differences remains challenging.

To address this issue, we have developed a cell population identification method (ILoReg) that takes an alternative approach to dimensionality reduction by means of feature extraction. At the core of ILoReg lies a new clustering algorithm, iterative clustering projection (ICP), which transforms a gene expression matrix into a probability matrix containing probabilities of each cell belonging to *k* clusters. These continuous cluster probabilities provide a more practical representation of the clustering than discrete cluster labels, as they can be handled like extracted features, and they are then utilized in the consensus clustering approach that combines multiple randomly subsampled ICP solutions into a consensus solution. The consensus approach acts as a noise-reducing step prior to hierarchical clustering and visualization by nonlinear dimensionality reduction, such as t-distributed stochastic neighbor embedding (t-SNE)^3^ or uniform manifold approximation and projection (UMAP)^4^. We have implemented this method as a user-friendly R package, ILoReg (https://github.com/elolab/iloreg), and demonstrate that it can greatly aid the identification of cell populations with subtle transcriptomic differences by increasing the cell population identification resolution of both clustering and visualization.

ICP is a clustering algorithm (Fig. 1a, **Supplementary Fig. 1** and **Methods**) that iteratively seeks a clustering of size *k* that maximizes the clustering similarity between the clustering and its projection by logistic regression, measured by the adjusted Rand Index (ARI). A particularly distinctive feature of ICP is the integration of feature selection with clustering by L1-regularized logistic regression, which at each step of the iteration automatically selects and weights genes by their relevance in predicting the current clustering. Since ICP generates different clusterings with different random seeds (**Supplementary Fig. 2**), ILoReg finally applies a consensus approach (Fig. 1b and **Methods**) to obtain a more accurate and robust solution, which is not constrained to the initial number of clusters *k* (**Supplementary Fig. 3**). ICP is run *L* times and the resulting cluster probability matrices are merged into a joint probability matrix, which is transformed into a lower dimension by principal component analysis (PCA). The PCA-transformed data matrix is clustered hierarchically using the Ward’s method and visualized using a nonlinear dimensionality reduction method, such as t-SNE or UMAP. To estimate the optimal number of clusters, ILoReg uses the silhouette method^5^.

**Figure 1.**
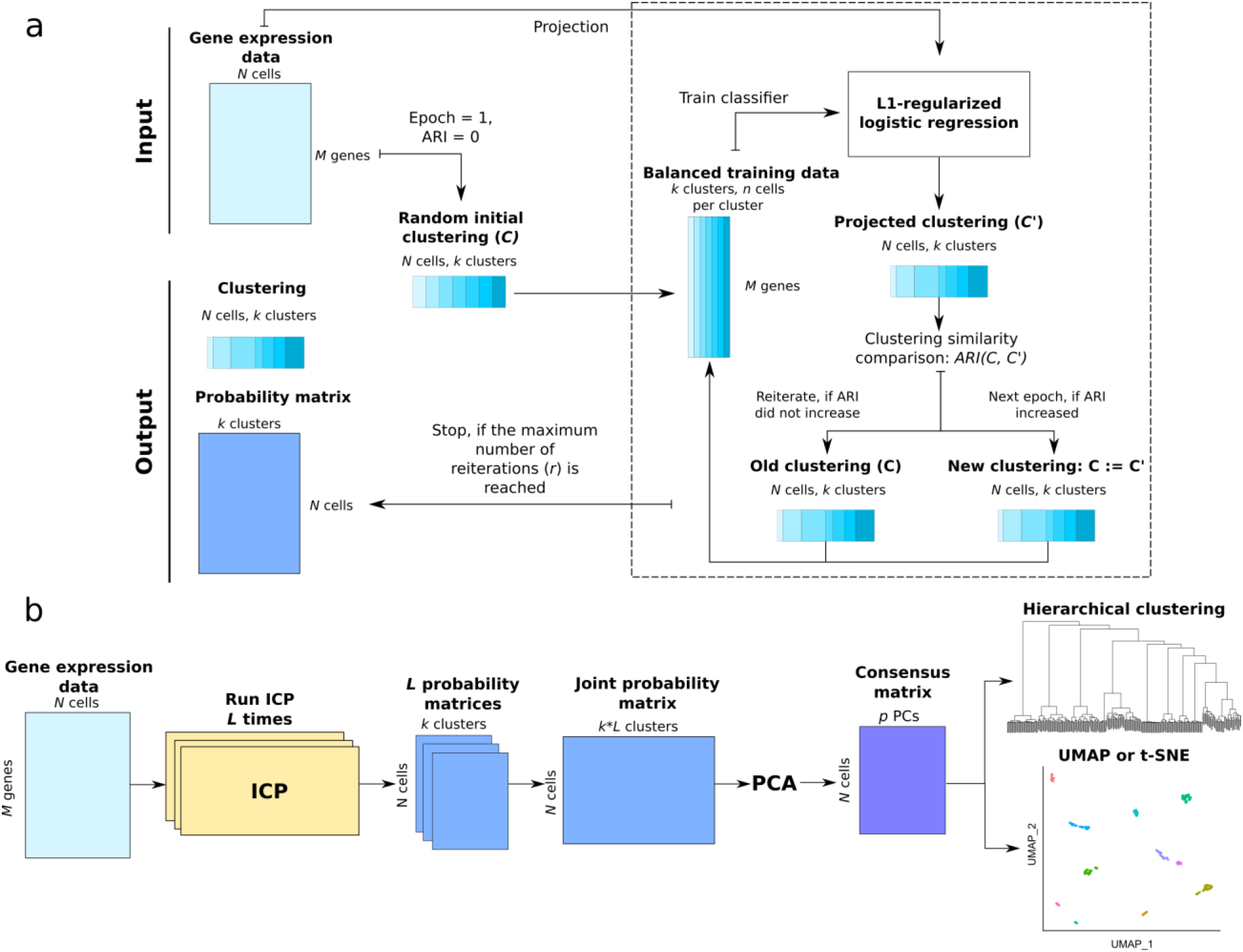
Overview of ILoReg. (**a**) Schematic of the iterative clustering projection (ICP) clustering algorithm. (**b**) Schematic of the ILoReg consensus approach for cell population identification.

We benchmarked ILoReg against four other clustering methods^6–9^, Seurat, SC3, CIDR and RaceID3, each functioning on a largely different principle (**Supplementary Table 1**), using eleven gold (Pollen) or silver (Baron and van Galen data) standard datasets from three publicly available studies^10–12^ (**Methods** and **Supplementary Table 2**). Although all the algorithms were able to find a number of clusters that was close to what the authors of the original studies reported (**Supplementary Fig. 4**), comparison by ARI between the inferred and original clusterings revealed considerable inaccuracies (Fig. 2a). ILoReg performed generally well regardless of the sample size, whereas CIDR and RaceID3 performed worse on average. In contrast to the other methods, SC3 tended to overestimate for larger datasets (e.g. Baron) and it was more accurate with smaller datasets (e.g. Pollen). Seurat performed consistently across the datasets, but for most datasets worse than ILoReg. In two of the datasets from the van Galen study (BM5-34p and BM5-34p38n), none of the methods were able to achieve even moderate accuracy for them. These datasets were sorted by flow cytometry and had highly unbalanced cluster labels in the original study, whereas in our comparison the inferred clusterings were more uniformly distributed, explaining the discrepancy.

**Figure 2.**
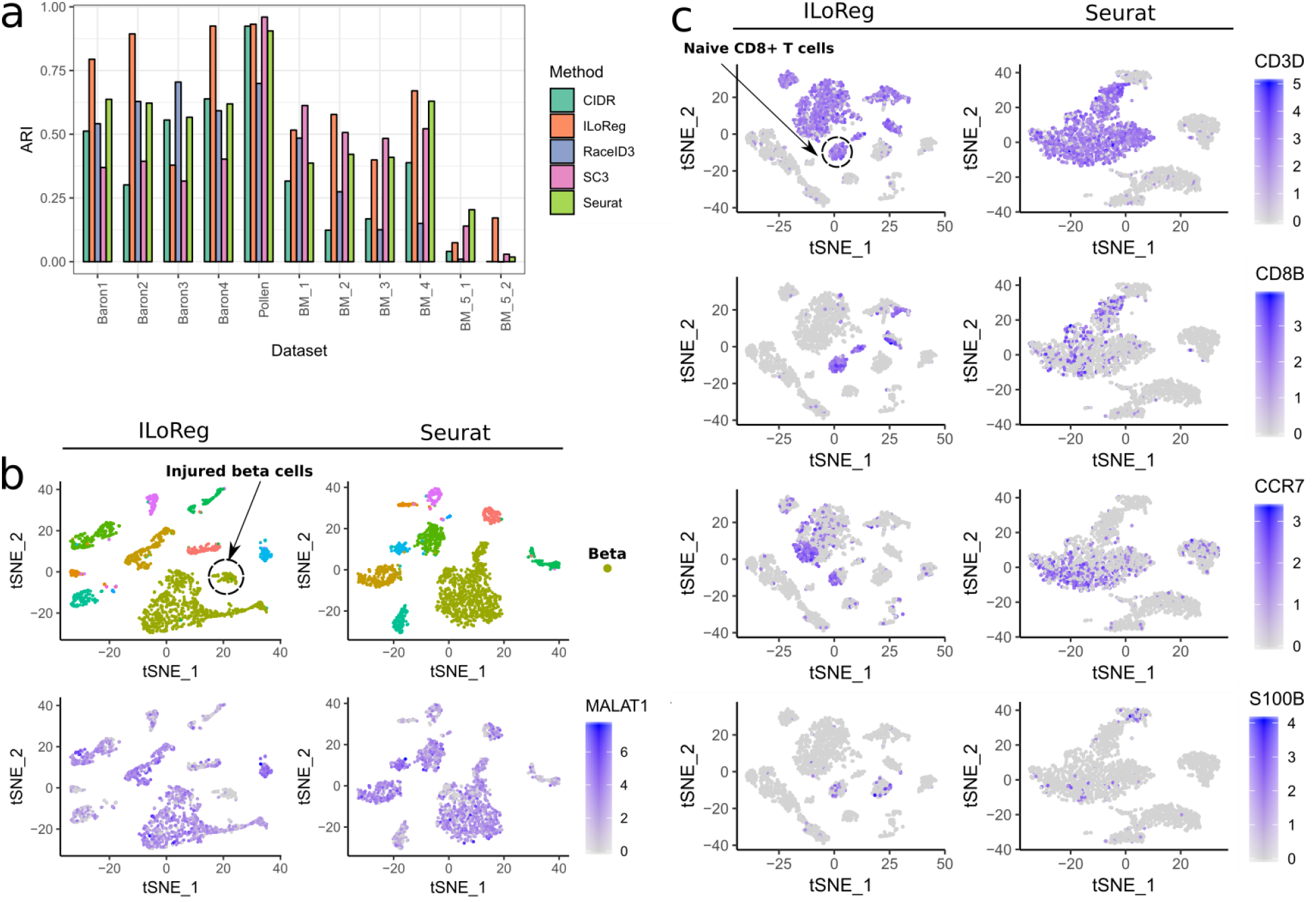
Benchmarking and identification of cell populations with subtle transcriptomic differences. **(a)** Benchmarking of ILoReg against four other scRNA-seq clustering methods: Seurat, SC3, CIDR and RaceID3. To compare the methods, the adjusted Rand index (ARI) was calculated in eleven datasets between the inferred clustering of each method and the reference clustering from the original study. The estimated numbers of clusters with each method were used; the estimates are provided in **Supplementary Figure 4**. The datasets with the “BM” prefix are from the van Galen study. **(b)** Comparison of *t*-distributed stochastic neighbor embedding (t-SNE) plots generated by ILoReg and Seurat for the Baron1 dataset, highlighting the expression levels of *MALAT1* that differentiates injured from healthy beta cells. The cells were colored by the reference labels from the original study. **(c)** Comparison of t-SNE plots generated by ILoReg and Seurat for the pbmc3k dataset, highlighting the expression levels of four genes (*CD3D*, *CD8B*, *CCR7*, *S100B*) that differentiate naive CD8+ T cells. All the analyses were carried out using the default parameter values of Seurat and ILoReg (*k = 15, C = 0.3, d = 0.3, r = 5, p = 50, L = 200*).

To demonstrate the ability of ILoReg to identify cell populations with subtle differences, we investigated in more detail clusters found from two datasets: a human pancreas dataset (Baron1) and a human peripheral blood mononuclear cell (PBMC) dataset (pbmc3k). From the Baron1 dataset, ILoReg identified a subpopulation of beta cells (Fig. 2b) with *MALAT1* downregulated, (Wilcoxon rank-sum test, adjusted *P* < 0.01, log2 FC~ −1.5). *MALAT1* is a gene that inhibits apoptosis and has been found to be negatively correlated with post-isolation islet cell death^13^. In line with this, functional analysis (**Methods**) of the differentially expressed genes between the two beta cell populations revealed a process that has been previously linked to beta cell destruction, i.e. endoplasmic reticulum stress (**Supplementary File 1**), further indicating the cluster indeed comprises injured beta cells^14^. The beta cells in the t-SNE representation of Seurat, however, are clustered more densely. Although an outer part of the whole beta cell cluster seems to contain *MALAT1*-beta cells, there is no clear separation between the *MALAT1*- and *MALAT1*+ cell populations.

In the pbmc3k dataset, a comparison between the five benchmarked methods (**Supplementary Fig. 5**) showed that the two-dimensional visualizations of all the four other methods (Seurat, SC3, CIDR and RaceID3) were similar, containing three main clusters: (1) T cells and NK cells; 2) dendritic cells and monocytes; 3) B cells. On the contrary, multiple distinct subpopulations are clearly visible within each main cell type in the t-SNE representation of ILoReg. Interestingly, unlike the other methods, ILoReg identified a cluster that expressed *CD3D, CD8B, CCR7* and *S100B* genes (Fig. 2c), corresponding to naive CD8+ T cells^15^. Overall, based on the expression of *CD8B* the separation of CD8- and CD8+ T cells by ILoReg was distinctly more precise. A more comprehensive analysis (**Supplementary Results 1**) revealed further cell populations that are in agreement with past studies, such as CD56+ and CD56++ NK cells^16^, naive and memory B cells with lambda or kappa light chain^17–19^, as well as rare platelets, which typically constitute less than 1% of PBMCs^20^.

The choice of the parameter values in ILoReg can have a considerable effect on the identifiable cell populations and they can be used to fine-tune the clustering to identify different cell populations. The effect is greater the higher the transcriptomic similarity between the populations is. The default parameter values were chosen so that biologically meaningful subpopulations with subtle expression differences are identified. We studied the effect of each of the parameters in detail and provide guidelines on how to select appropriate parameter values (**Methods**).

The most computationally intensive part of ILoReg is running ICP *L* times (default *L* = 200). To accelerate it, the R package supports computing the ICP runs in parallel. Using the default parameter values and 12 threads, the run times for ~3,000 cells and ~20,000 cells were ~1 hr and ~10 hr, respectively (**Methods** and **Supplementary Fig. 6**). It should be noted, however, that although the run time of a single workflow was relatively long, the workflow is generally simple and the number of consensus clusters *K* is very fast to change (~1 s with ~3,000 cells). By contrast, with SC3 the user must repeat the clustering for each number of clusters *k* separately, thus roughly multiplying the run time by the number of different *k* values. Similarly, with Seurat the user must repeat the graph-based clustering with different resolution values without knowing how many clusters a resolution value gives. Another considerable benefit of ILoReg is its ability to find subpopulations without the need of further sub-clustering, therefore considerably simplifying the analysis workflow and saving further time.

To conclude, we have developed a new cell population identification method, ILoReg, which enables high-resolution identification of cell subpopulations from scRNA-seq data by a consensus clustering approach. Remarkably, using the cluster probabilities in feature extraction increased the resolution of nonlinear dimensionality reduction methods, such as t-SNE or UMAP, compared to current state-of-the-art methods that preprocess data by selecting a set of highly variable genes. In particular, our results demonstrate that ILoReg can greatly aid the identification of cell populations with subtle transcriptomic differences.

## Methods

### Iterative clustering projection

As a basis of ILoReg, we first introduce a new clustering algorithm, iterative clustering projection (ICP), that utilizes random down- and oversampling, supervised learning and clustering comparison to iteratively cluster the data (Fig. 1a and **Supplementary Fig. 1**). Specifically, the objective of ICP is to seek a clustering *C* = {*C*_1_, …, *C*_*k*_} with *k* clusters that maximizes the adjusted Rand index (ARI) between *C* and its projection *C*′ by logistic regression:

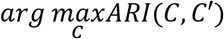

In the following, we describe the four steps of the algorithm.

1. *Initialization*. Given a normalized gene expression matrix, *X* = [*x*_1_, …, *x*_*N*_]^*T*^, where *N* is the number of cells and *x*_*i*_ is the transcriptional profile of the *i*th cell across *M* genes that are expressed in at least one of the *N* cells, the algorithm first splits the cells randomly into *k* clusters *C*_*t*_ = {*C*_1,*t*_, …, *C*_*k*,*t*_}using a uniform probability distribution, where *t* = 1.
2. *Creating balanced training data*. To form a balanced training dataset *X*_*t*_ and training labels *Y*_*t*_ = {*Y*_1,*t*_, …, *Y*_*k*,*t*_} with an equal number of cells *n* in each cluster of *Y*_*t*_, down- and oversampling of *C*_*t*_ and *X*are carried out. If a cluster contains more than *n* cells, then its cells are randomly downsampled by selecting *n* cells from the cluster without replacement. If a cluster has fewer than *n* cells, then its cells are oversampled with replacement. Because scRNA-seq datasets come at very different sample sizes, *n* is determined by *n* = ⌈*Nd*/*k*⌉, where *d* ∈ (0,1) and ⌈⌉denotes the ceiling function.
3. *Classifier training and projection*. An L1-regularized logistic regression classifier is trained on the training data *X*_*t*_ and labels *Y*_*t*_using the LIBLINEAR library^21^. *X* is projected onto itself with the classifier, i.e. the cluster of each of the *N* cells is predicted with *X* as input data, which yields the projected clustering*C*′_*t*_ = {*C*′_1,*t*_, …, *C*′_*k*,*t*_} and the probability matrix *P*_*t*_ = [*p*_1,*t*_, …, *p*_*N*,*t*_]^*T*^, where *p*_*i*,*t*_is a real vector containing the probabilities of the *i*th cell belonging to the *k* clusters. The objective function of L1-regularized logistic regression is

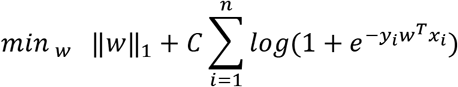

where *w* is the model weight vector, *n* the number of training samples in the *k*th cluster, *y*_*i*_ = {−1,1} and ‖‖_1_ the 1-norm. The constant *C* > 0 determines the trade-off between regularization and correct classification, a lower value selecting fewer genes. To perform multiclass classification, the LIBLINEAR library uses the one-vs-rest scheme, in which *k* binary classifiers are trained using the samples belonging to one of the *k* clusters as positive samples and all other samples from the *k* − 1 clusters as negatives.
4. *Clustering comparison*. The similarity between clusterings *C*_*t*_ and *C*′_*t*_ is measured by ARI^22^:

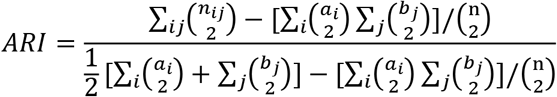 Here *n* is the total number of cells, *n*_*ij*_ is the number of overlapping cells in clusters *i* and *j* from *C*_*t*_ and *C*′_*t*_ respectively, *a*_*i*_ and *b*_*j*_are the total number of cells in clusters *i* and *j* from*C*_*t*_ and *C*′_*t*_ respectively. If ARI increases from its previous value (initialized to 0 with *t* = 1), then *C*_*t*+1_is set to *C*′_*t*_ and steps 2, 3 and 4 are repeated for *C*_*t*+1_in the next epoch. If ARI does not increase, the steps 2, 3 and 4 are repeated for *C*_*t*_ until the maximum number of reiterations *r* is reached. At the start of every new epoch, the number of reiterations is set to 0. After the last reiteration, *P*_*t*_ and *C*′_*t*_ are returned as output, where *t* is the last epoch.

To understand how the parameters of ICP (*d*, *C*, *k* and *r*) affect the identification of cell populations and constrain their values, we investigated adjusting their values with the pbmc3k dataset and studied how that affected cell populations found by ILoReg (**Supplementary Fig. 7-10**).

The parameter *d* enables controlling the diversity of the consensus solution. While a higher *d* increased the projection accuracy (**Supplementary Fig. 11a**), it also reduced the average similarity of different ICP runs (**Supplementary Fig. 11b**), which in turn induced less variation into the consensus solution (**Supplementary Fig. 11c**). This is consistent with the t-SNE visualizations generated using different values of *d* (**Supplementary Fig. 7**), containing fewer visible subpopulations of T cells for higher *d* values. Additionally, since with *d* = 0.1 and *C = 0.1* the number of clusters *k* started to randomly decrease during the iteration due to the clustering becoming unbalanced, using *d* smaller than 0.2 is not recommended.

The parameter *C* controls the stringency of the feature selection through L1-regularization, a lower value selecting fewer genes into the logistic regression model. With a lower *C* ICP achieved a higher projection accuracy at the final epoch of ICP (**Supplementary Fig. 11a**). From *C* = 0.3 to *C* = 1.0 the t-SNE plots had similar shapes for the populations that expressed the three T cell marker genes (*CD8B, CCR7, TNFRSF4*), but at *C* = 0.1 the result differed significantly more (**Supplementary Fig. 8**).

The number of initial clusters *k* determines the dimensionality of the cluster probability matrix, and intuitively a higher *k* increased the cell population diversity of the result (**Supplementary Fig. 9**). With *k* = 5 and *k* = 10 CD4+ T cells were distinctly grouped into sharp clusters based on the expression of *CCR7*. Although this kind of extreme differentiation resolution may be useful, many real cell types underlying the tissue may not be separable by on-off expression of a single gene. For example, both naive and memory T cells have been shown to express *CCR7*, but its expression level is, on average, higher in naive T cells^23^. Therefore, using *k* = 15 provided an embedding that was more consistent with past studies.

The maximum number of reiterations *r* enables controlling the projection accuracy of ICP without the need to adjust the values of *k*, *d*, and *C*. Based on the overall high similarity of the t-SNE representations that were acquired using different values of *r* (**Supplementary Fig. 10**), *r* should be of least concern when tuning the parameters.

### Consensus clustering method

A consensus method (Fig. 1b) is used to obtain a more accurate and robust clustering of the cells than the clusterings obtained by the individual ICP runs (**Supplementary Fig. 3**), which is also not constrained to the number of initial clusters *k*. The ICP algorithm is run *L* times with different random seeds and their probability matrices are merged to create the joint probability matrix, *P* = [*P*_1_, …, *P*_*L*_]. The dimensionality of the data is reduced with principal component analysis (PCA) by performing eigendecomposition of the cross-product of the centered *P*using the RSpectra R package. Finally, the *N* × *p*dimensional consensus matrix is clustered efficiently (*O*(*Np*)) using hierarchical clustering with the Ward’s method from the fastcluster R package^24^. The tree dendrogram from the hierarchical clustering is cut into *K* clusters with the dendextend R package^25^. The optimal number of clusters can be determined automatically by the silhouette method from the cluster R package.

Since there were only minor differences between the t-SNE representations computed with *p* = 50 and *p* = 100 (**Supplementary Fig. 12**), and a common strategy among methods that apply PCA is to select rather too many components than too few (e.g. the Rtsne R package), we decided to use 50 as the default value of *p*. The ILoReg R package also provides a function for drawing an elbow plot to help the user in selecting the number of principal components. Additionally, we investigated the impact of the parameters *L* and *k*, suggesting (**Supplementary Fig. 3**) that regardless of which of the different values of *k* and *L* were used, the consensus method was able to find a clustering that was accurate when measured by ARI. The consensus clustering stabilized when *L* was between 10 and 50, being also in agreement with the t-SNE plots computed with different values of *L* (**Supplementary Fig. 13**). However, this is likely to be dataset-specific and therefore to be safe, we set the default value to 200.

### Visualization

ILoReg supports visualization using two popular nonlinear dimensionality reduction methods: t-distributed stochastic neighbor embedding (t-SNE) from the Rtsne R package and uniform manifold approximation and projection (UMAP) from the umap R package. The *p*-dimensional PCA-transformed matrix is used as input in both of the methods.

### Benchmarking

To benchmark ILoReg against other scRNA-seq clustering methods, we considered four state-of-the-art methods: Seurat^6^, SC3^7^, CIDR^8^ and RaceID3^9^. Details of the methods are listed in **Supplementary Table 1**. We carried out the benchmarking assuming that the true number of clusters is unknown, and therefore, used the default parameter values with each method. With RaceID3 we used the initial clustering by *k*-medoids without the subsequent outlier detection step that adds further clusters. Seurat is the only method without automated estimation of *k*, and we therefore used the number of clusters provided by the default resolution value 0.8. To measure clustering accuracy, we used ARI between the ground truth and inferred clusterings. Additionally, we compared the estimated and ground truth values of *k*.

For benchmarking we selected three public scRNA-seq datasets, in which each cell has been categorised by the authors of the original publication. The benchmarking datasets that we used are listed in **Supplementary Table 2**. The Baron and Galen datasets are silver standard datasets, i.e. their clusters were identified by the authors based on gene markers, with several replicates. The Pollen dataset is a gold standard dataset with information on which cell line each cell originated from. Before clustering we removed spike-ins from the Pollen dataset. The normalization of each dataset was performed using the same method that was given in the original study.

### Preprocessing of the pbmc3k dataset

The raw FASTQ reads of the pbmc3k dataset were downloaded from the public database of the 10X Genomics company (https://support.10xgenomics.com/single-cell-gene-expression/datasets) and the preprocessing was performed using Cell Ranger v2.2.0 and the GRCh38.p12 human reference genome. The unique molecular identifier (UMI) counts were normalized using the LogNormalize method from the Seurat R package.

### Functional analysis of the Baron1 dataset

To identify enriched biological pathways among the differentially expressed genes between the healthy and injured beta cell populations from the Baron1 dataset, we used the Metascape web tool^27^.

### Run time and memory usage

The run time and maximal resident set size (RSS) of the five benchmarked methods were measured using two PBMC datasets: pbmc3k (~ 3k cells) and a subset of the fresh_68k_pbmc_donor_a dataset (20k cells) on a cluster node with CentOS Linux 7 operating system, 12-core 2.66 GHz Intel Xeon X5650 processor and 96 GB 1066 MHz DDR3 of RAM (**Supplementary Fig. 6**). The workflow steps that were included in this comparison were dimensionality reduction, clustering and estimating the optimal number of clusters. In contrast to the other methods, changing the number of clusters *k* with SC3 can be time-consuming due to the computational bottleneck step involving *k*-means clustering. Since in practice the user needs to run the consensus clustering with a range of different *k* values, we adjusted the SC3 workflow to use *k* values ranging from 2 to 50. 12 threads were used with the methods that support parallel computing (SC3 and ILoReg).

### Differential expression analysis

The ILoReg R package provides user-friendly functions that enable identification of gene markers for clusters and visualization of gene expression across cells and between clusters (**Supplementary Fig. 5**). The current implementation of ILoReg supports two functions for gene marker identification. The first function, *FindAllGeneMarkers*, allows simultaneous identification of gene markers for all K clusters. Differential expression analysis is performed using the one-vs-rest scheme, in which cells from each cluster are compared against the rest of the cells. To accelerate the analysis, the user can apply filters to remove genes that are less likely to be good marker genes or downsample cells. The differential expression analysis uses the Wilcoxon rank-sum test to calculate a p-value representing the statistical significance of a gene. The p-value adjustment for multiple comparisons is carried out using the Bonferroni method. A second function, *FindGeneMarkers*, enables comparison between any two arbitrary sets of clusters.

### Datasets

The Galen dataset was downloaded from the NCBI’s GEO database (GSE116256). The PBMC and pan T cell datasets were downloaded from the public database of the 10X Genomics company (https://support.10xgenomics.com/single-cell-gene-expression/datasets). Baron and Pollen datasets were downloaded from the public database of the Hemberg lab (https://hemberg-lab.github.io/scRNA.seq.datasets/).

### Software availability

ILoReg is available as an R package at https://github.com/elolab/iloreg.

The version of ILoReg used for generating the results in this manuscript can be downloaded from GitHub using the reference ID “85196be6”.

All analyses in this study were performed using R (version 3.6.0) and the following versions of the R packages were used:

- SC3 (version 1.12.0)
- Seurat (version 3.0.0)
- CIDR (version 0.1.5)
- RaceID (version 0.1.3)
- Rtsne (version 0.15)
- umap (version 0.2.0.0)
- cluster (version 2.0.8)
- RSpectra (version 0.14-0)
- aricode (version 0.1.1)
- DescTools (version 0.99.28)
- LiblineaR (version 2.10-8)
- dendextend (version 1.10.0)
- cowplot (version 0.9.4)
- ggplot2 (version 3.1.1)
- SparseM (version 1.77)
- doParallel (version 1.0.14)
- foreach (version 1.4.4)
- pheatmap (version 1.0.12)

## Acknowledgements

Prof. Elo reports grants from the European Research Council ERC (677943), European Union’s Horizon 2020 research and innovation programme (675395), Academy of Finland (296801, 304995, 310561 and 314443), Juvenile Diabetes Research Foundation JDRF (2-2013-32), and Sigrid Juselius Foundation, during the conduct of the study. Our research is also supported by University of Turku, Åbo Akademi University, Turku Graduate School (UTUGS), Biocenter Finland, and ELIXIR Finland. The authors thank the Elo lab for fruitful discussions and comments on the manuscript.

## Author contributions

J.S. and L.L.E conceived the study; J.S. invented and implemented the method and designed the study; J.S. and S.J. developed the computational framework; S.J., M.S.V. and L.L.E. supervised the research; and J.S. wrote the manuscript with the help of the other authors.

## Competing interests

The authors declare no competing interests.

