## Supplementary Information for "ILoReg enables high-resolution cell population identification from single-cell RNA-seq data"

#### Supplementary Figures

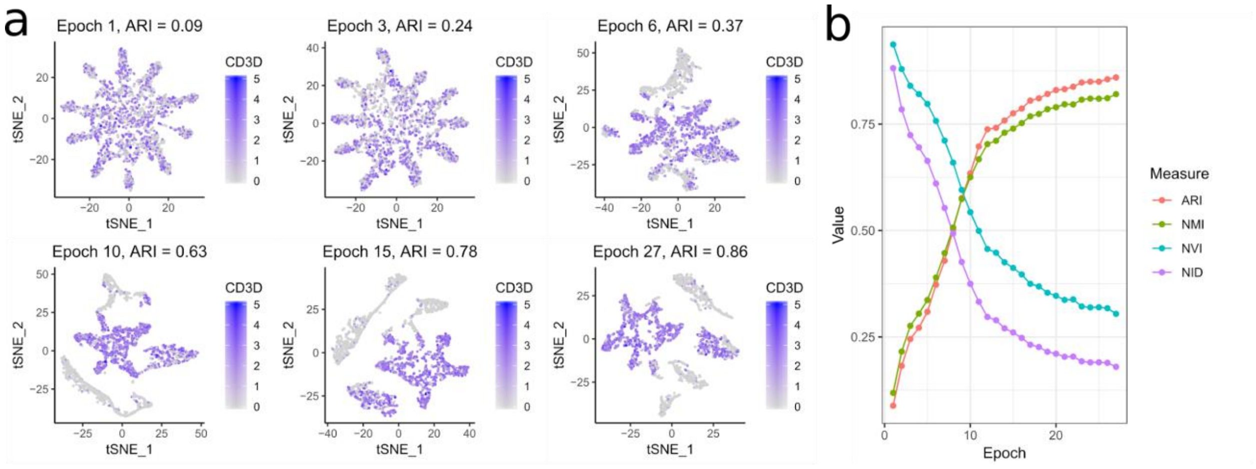


**Supplementary Figure 1**

Convergence of the ICP algorithm

**(a)** *t*-distributed stochastic neighbor embedding (t-SNE) transformations of the *N* x *k* -dimensional probability matrix at different epochs of the iterative clustering projection (ICP) clustering algorithm, where *N* is the total number of cells and *k* is the number of clusters. The expression levels of the T cell marker gene *CD3D* are highlighted. **(b)** Clustering comparison measures calculated between the clustering and its projection at every epoch: adjusted Rand index (ARI), normalized mutual information (NMI), normalized variation information (NVI) and normalized information distance (NID). ARI at the final epoch is denoted as the projection accuracy of ICP. See **Supplementary Figure 11** for an analysis on how the *d* and *C* parameters affected the projection accuracy of ICP.

In this example, we used the pbmc3k dataset with $k$ = 10, $C$ = 0.3, $d$ = 0.3 and $r$ = 5 parameter values. The clustering comparison measures in **(b)** were calculated using the aricode R package.

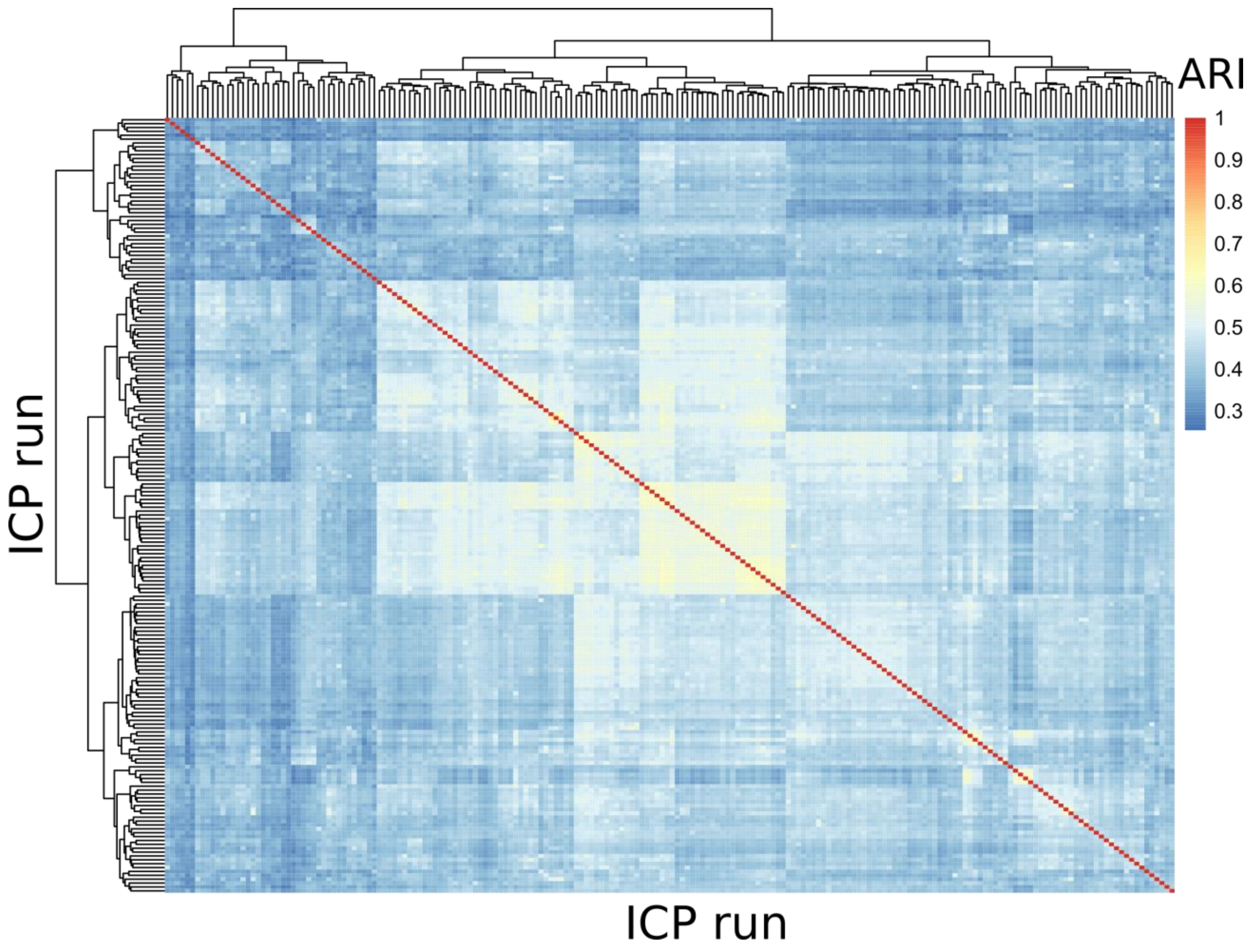


**Supplementary Figure 2**

Similarity between ICP runs

The iterative clustering projection (ICP) algorithm was run 200 times for the pbmc3k dataset using $k$ = 15, $C$ = 0.3, $d$ = 0.3 and $r$ = 5 parameter values. To measure similarity between the results of different ICP runs, the average pairwise adjusted Rand index (ARI) was calculated by taking the average of each row of the matrix. The heatmap and the hierarchical clustering were computed using the pheatmap R package with its default parameter values. See **Supplementary Figure 11** for an analysis on how the $d$ and $C$ parameters affected the similarity of ICP runs.

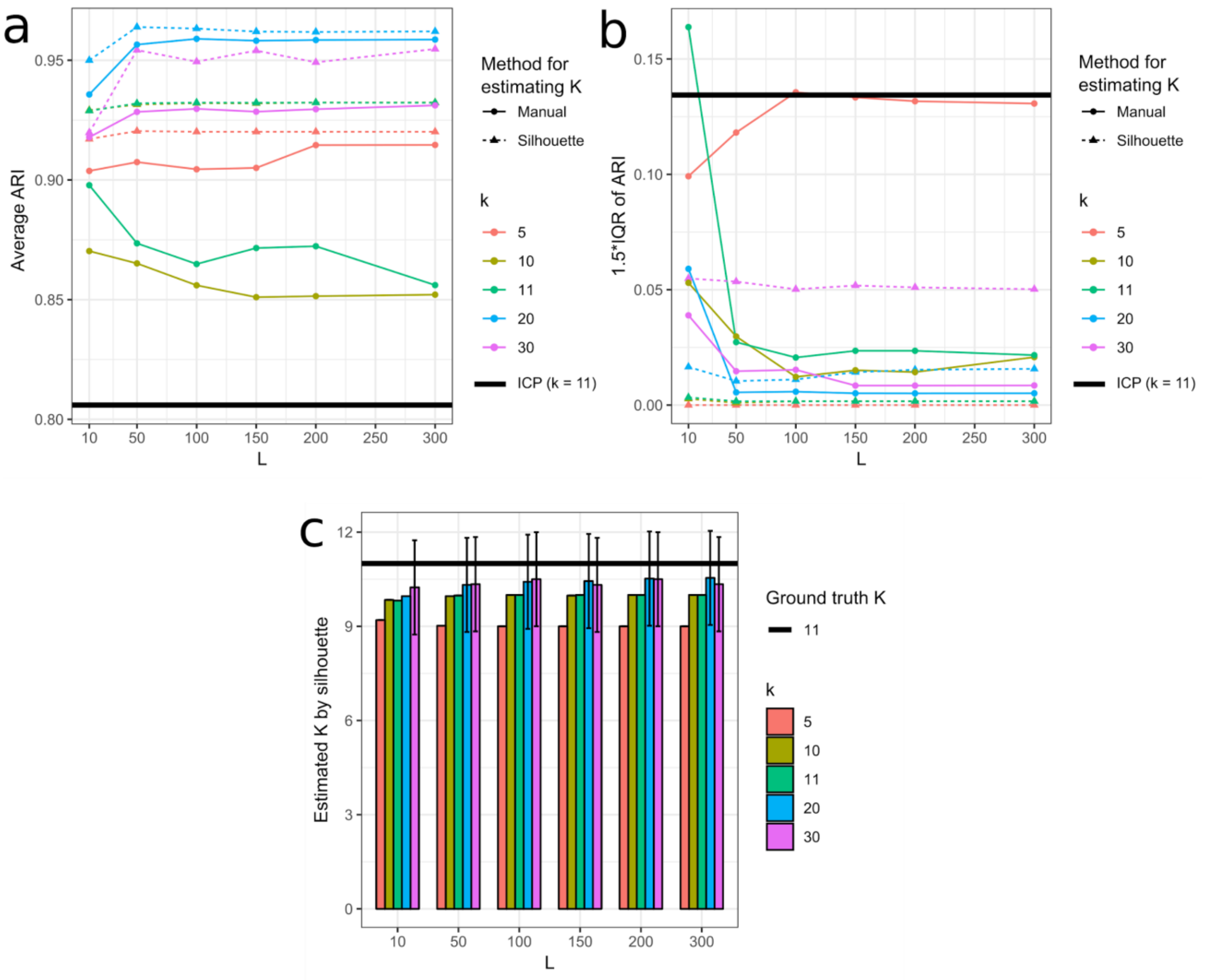


**Supplementary Figure 3**

Comparison of ICP and the ILoReg consensus method

**(a)** Coloured line plots showing the average adjusted Rand index (ARI) achieved by the ILoReg consensus clustering method using different values of the initial number of clusters (*k*) in iterative clustering projection (ICP) and the number of ICP runs (*L*). The average was taken over 50 different randomly initialized ILoReg consensus clustering solutions, where ARI was calculated between the inferred clustering and the reference clustering from the Pollen study. Additionally, two approaches for selecting the number of consensus clusters from the dendrogram (*K)* were compared: the silhouette method (Silhouette) and selecting the same number of clusters as in the reference clustering (Manual). The black horizontal line denotes the average ARI achieved by 500 individual ICP runs with the same number of clusters as in the reference clustering (*k* = 11). **(b)** Line plots depicting the variability of the results in **(a)**: 1.5* the interquartile range (IQR) of the ARI values. **(c)** The average number of estimated clusters by the silhouette method. The error bars depict 1.5*IQR of the estimated *K* from 50 different ILoReg consensus solutions.

The rest of the ILoReg parameters were fixed to *C* = 0.3 , d = 0.3, *r* = 5 and *p* = 50.

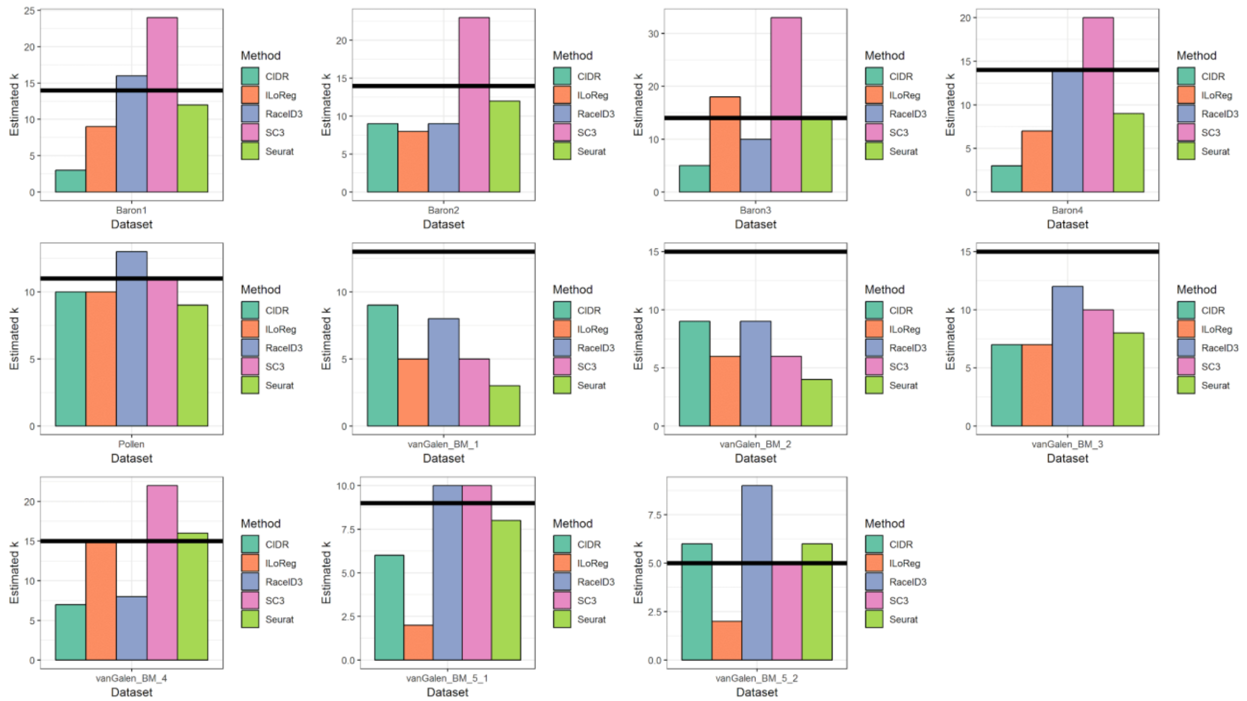


**Supplementary Figure 4**

Estimation of the number of clusters in benchmarking

Barplots showing the estimated number of clusters for each benchmarked method and dataset. The corresponding clusterings were used in the benchmarking (**Fig. 2**). The black horizontal line denotes the number of clusters found in the original studies.

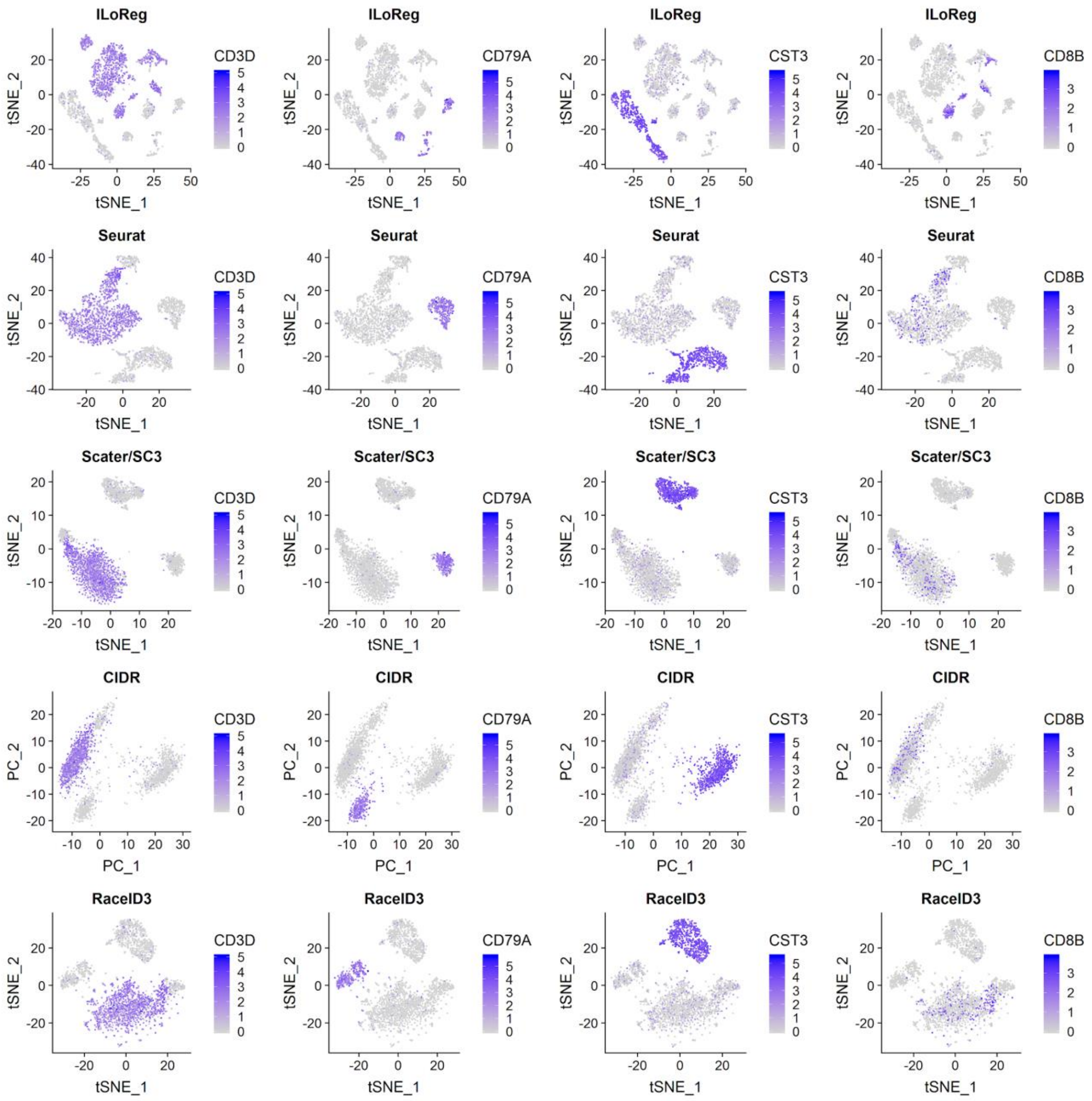


**Supplementary Figure 5**

Visualization comparison between the benchmarked methods

Expression levels of gene markers for major PBMC types, *CD3D* (a T cell marker), *CD79A* (a B cell marker), *CST3* (a marker for monocytes, dendritic cells and platelets) and *CD8B* (a CD8+ T cell marker) highlighted over two-dimensional visualizations of each benchmarked method.

SC3 is the only method without its own two-dimensional visualization function, but the scater R package[1] was used to perform the visualization, as recommended in the manual of the SC3 R package.

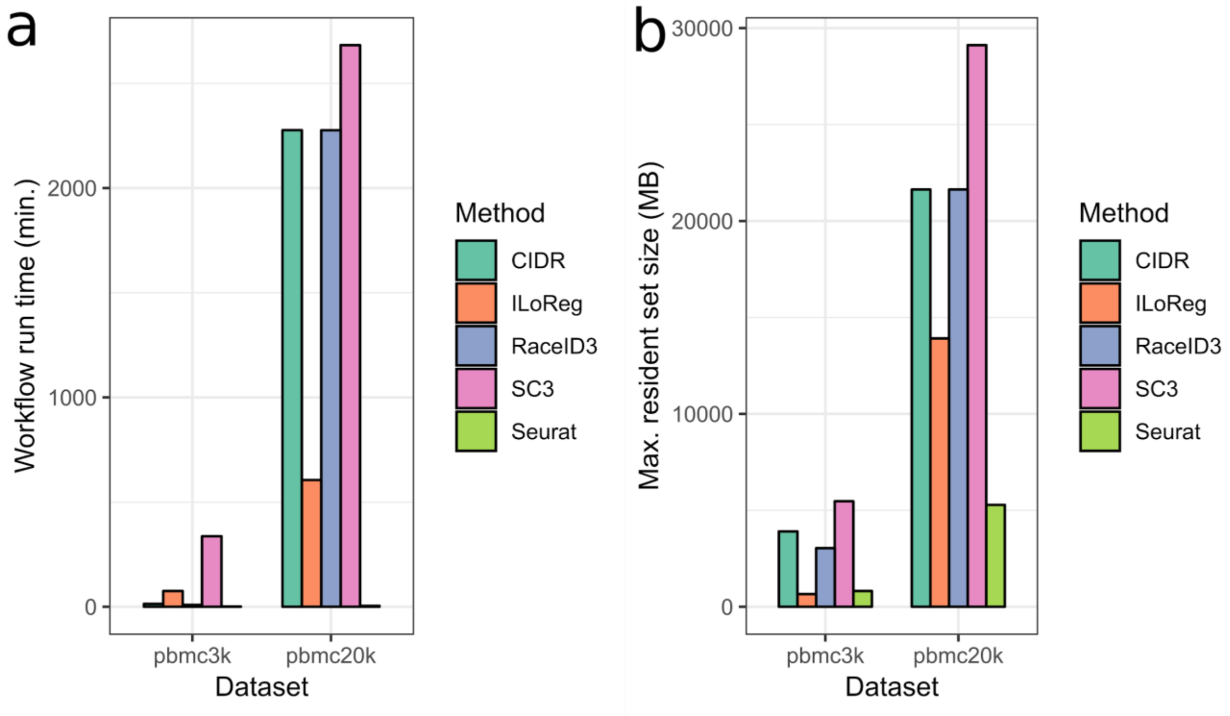


**Supplementary Figure 6**

Run time and memory usage comparison

**(a)** Run times for completing the analysis workflows of different methods. **(b)** Maximum amount of memory used at any time by any process during the workflows.

To clarify, pbmc3k and pbmc20k denote datasets with ~ 3,000 and 20,000 peripheral blood mononuclear cells (PBMCs), respectively. See **Methods** for further details of the run time and memory usage comparison.

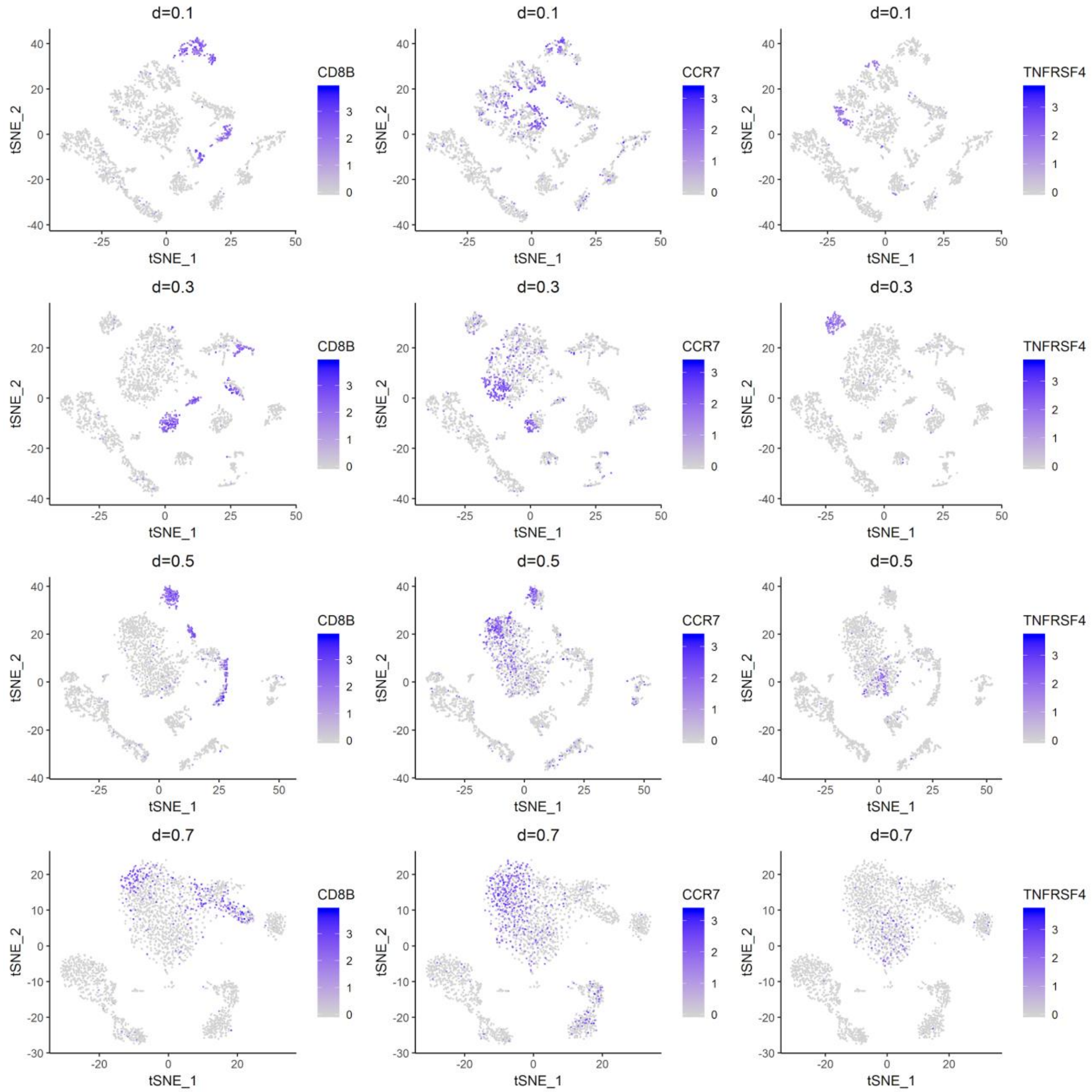


**Supplementary Figure 7**

Effect of the parameter *d* on the location of T cell subpopulations in t-SNE dimensionality reduction

*t*-distributed stochastic neighbor embedding (t-SNE) plots generated by ILoReg for the pbmc3k dataset with different values of the *d* parameter that controls the number of cells that are selected into the balanced training data in iterative clustering projection (ICP). The expression levels of three genes are highlighted: *CD8B* (CD8+ T cells), *CCR7* (naive T cells), *TNFRSF4* (activated CD4+ T cells). The other parameters were fixed: $C$ = 0.3, $k$ = 15, $r$ = 5, $L$ = 200, $p$ = 50.

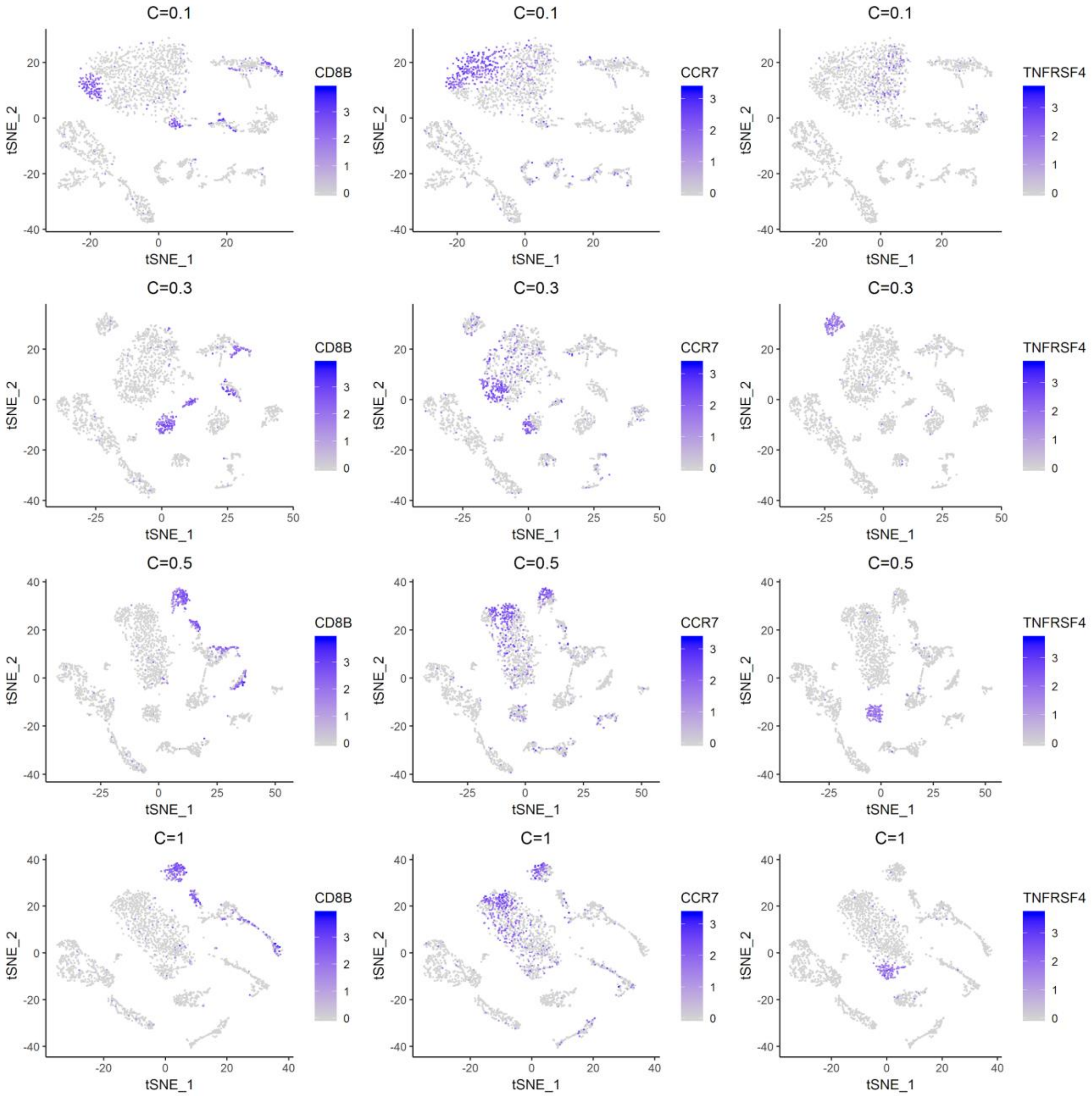


**Supplementary Figure 8**

Effect of the parameter *C* on the location of T cell subpopulations in t-SNE dimensionality reduction

*t*-distributed stochastic neighbor embedding (t-SNE) plots generated by ILoReg for the pbmc3k dataset with different values of the *C* parameter that controls the stringency of the L1-regularized feature selection in the logistic regression model of iterative clustering projection (ICP). The expression levels of three genes are highlighted: *CD8B* (CD8+ T cells), *CCR7* (naive T cells), *TNFRSF4* (activated CD4+ T cells). The other parameters were fixed: $d$ = 0.3, $k$ = 15, $r$ = 5, $L$ = 200, $p$ = 50.

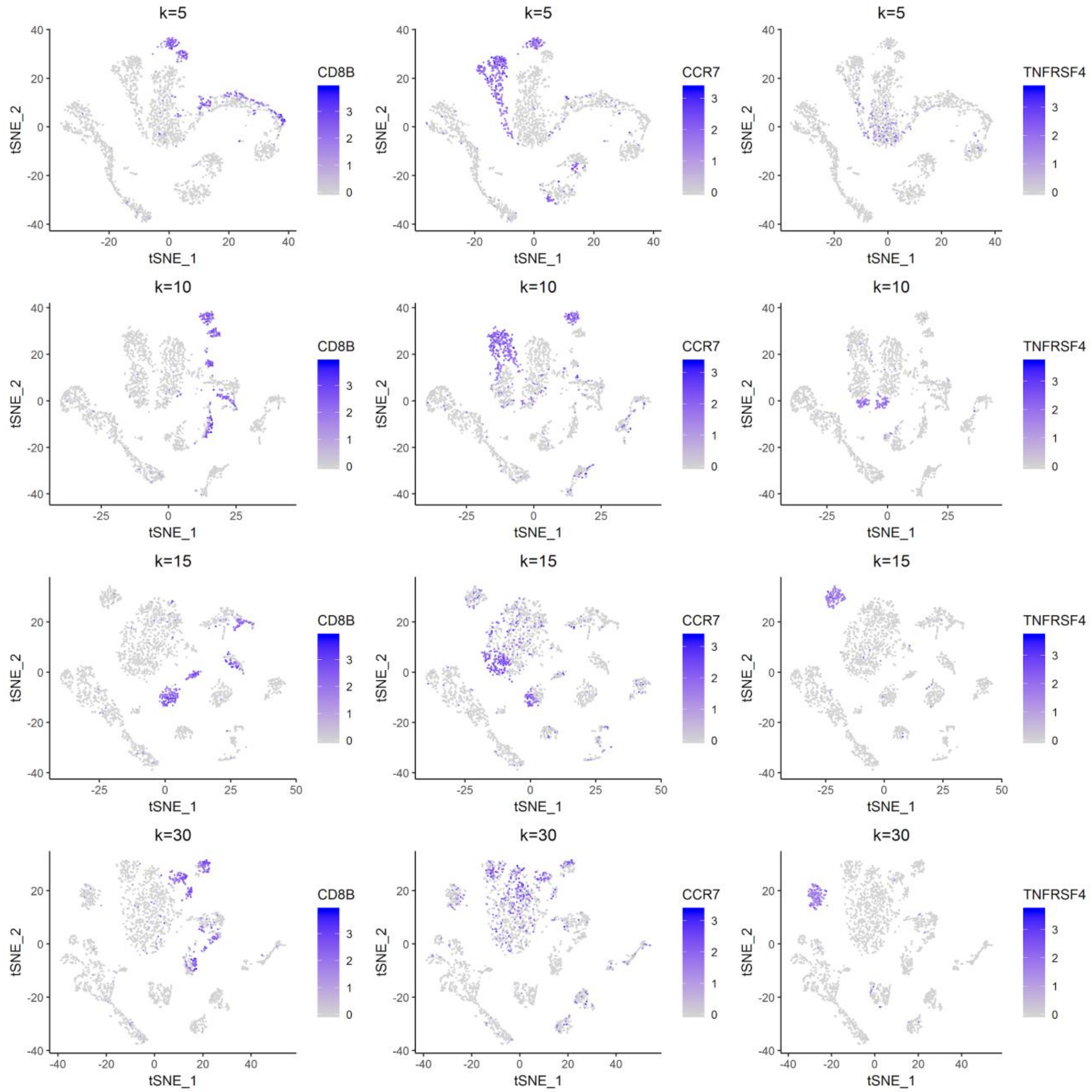


**Supplementary Figure 9**

Effect of the parameter *k* on the location of T cell subpopulations in t-SNE dimensionality reduction

*t*-distributed stochastic neighbor embedding (t-SNE) plots generated by ILoReg for the pbmc3k dataset with different values of the number of initial clusters (*k*) in iterative clustering projection (ICP). The expression levels of three genes are highlighted: *CD8B* (CD8+ T cells), *CCR7* (naive T cells), *TNFRSF4* (activated CD4+ T cells). The other parameters were fixed: $d$ = 0.3, $C = 0.3$, $r$ = 5, $L$ = 200, $p = 50$.

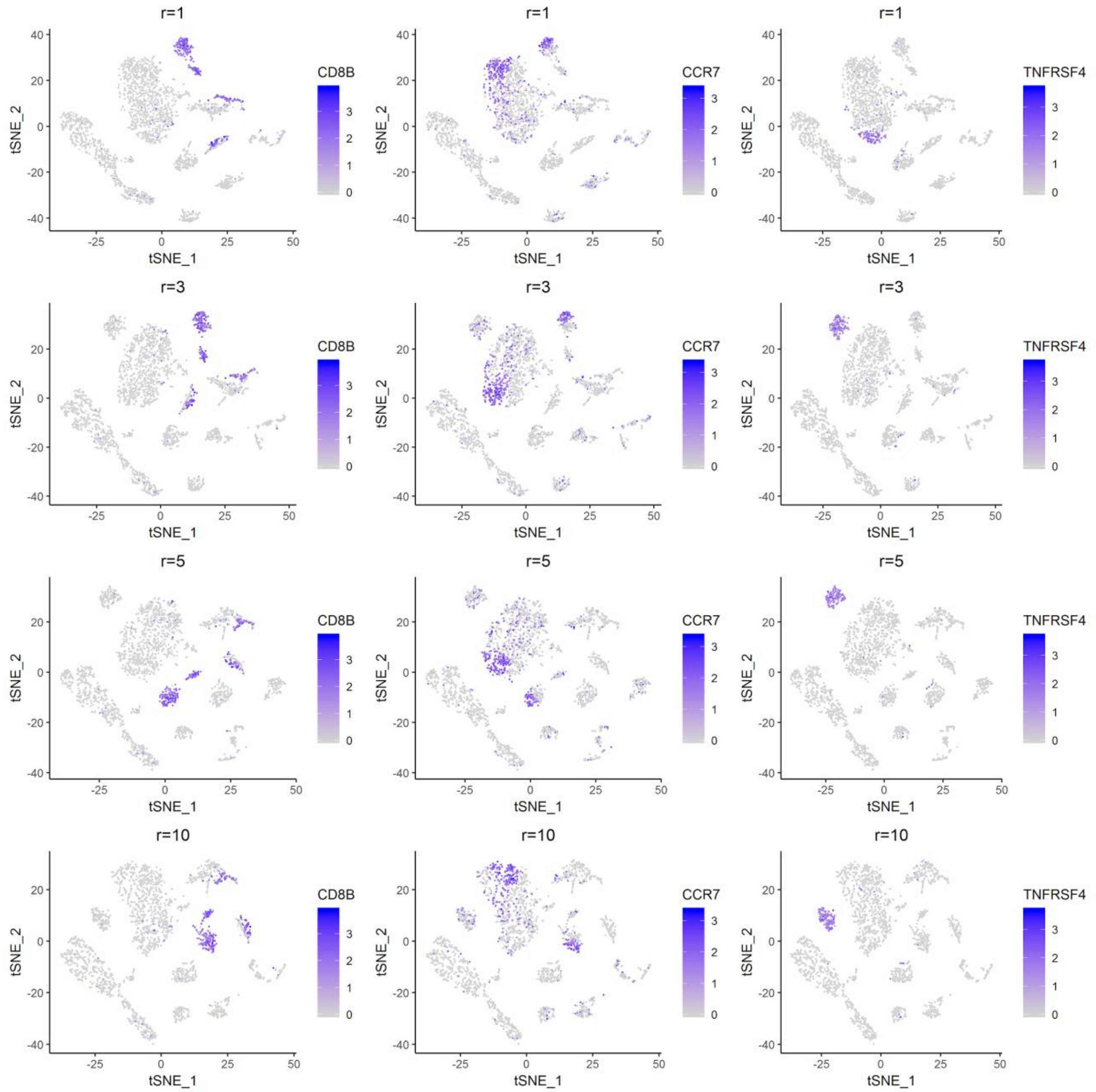


**Supplementary Figure 10**

Effect of the parameter *r* on the location of T cell subpopulations in t-SNE dimensionality reduction

*t*-distributed stochastic neighbor embedding (t-SNE) plots generated by ILoReg for the pbmc3k dataset with different values of the maximum number of reiterations (*r)* in iterative clustering projection (ICP). The expression levels of three genes are highlighted: *CD8B* (CD8+ T cells), *CCR7* (naive T cells), *TNFRSF4* (activated CD4+ T cells). The other parameters were fixed: *d* = 0.3*, C = 0.3, k* = 15*, L* = 200*, p = 50.*

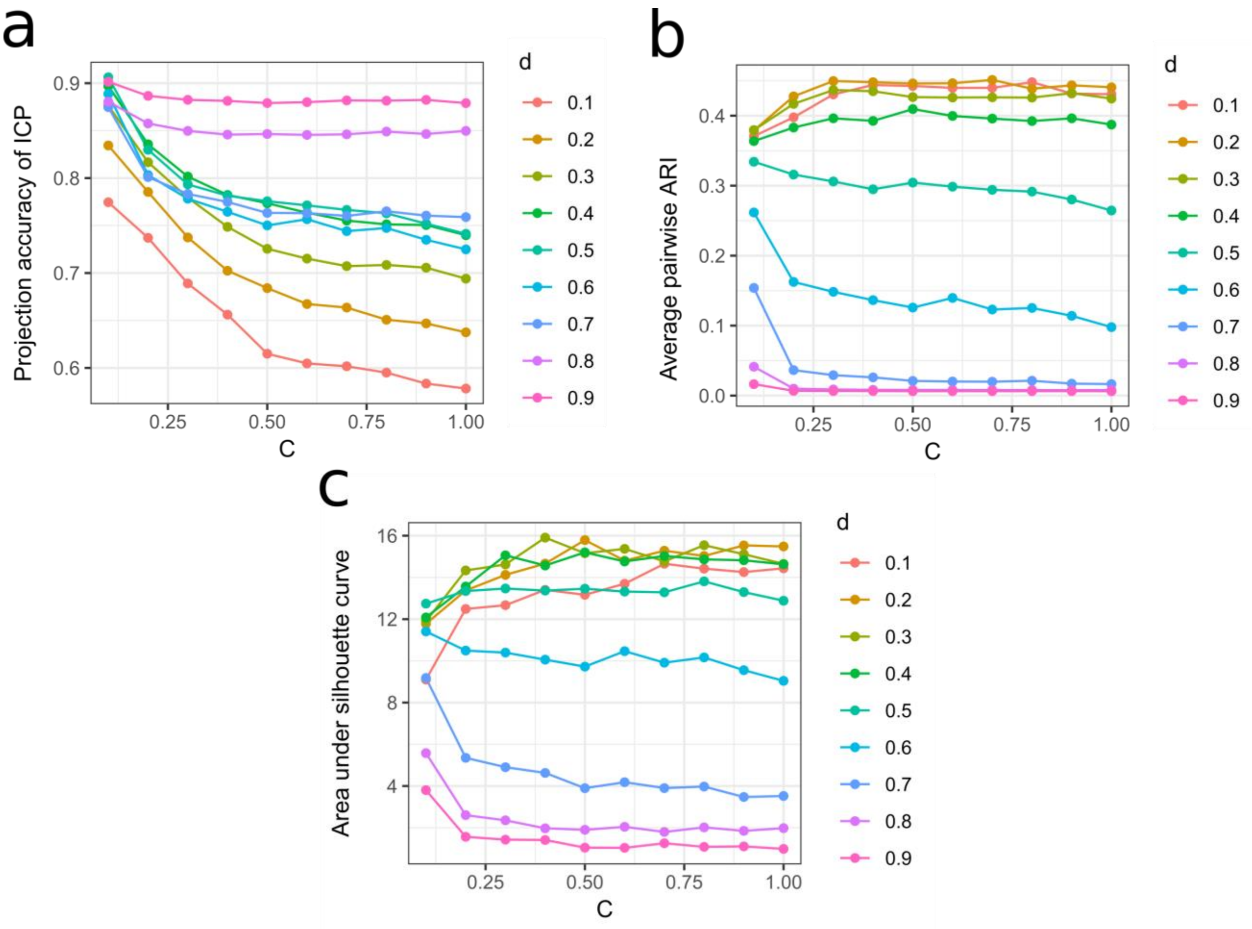


**Supplementary Figure 11**

Effect of the *C* and *d* parameters on the projection accuracy of ICP, the similarity of ICP runs and the area under the silhouette curve value of the ILoReg consensus clustering

**(a)** The projection accuracy (**Supplementary Fig. 1**) achieved by iterative clustering projection (ICP) using different values of the cost parameter in L1-regularized logistic regression (*C*) and the parameter controlling the number of cells per cluster in training data of logistic regression (*d*). **(b)** The average pairwise adjusted Rand index (ARI) (**Supplementary Fig. 2**) achieved using different values of the *C* and *d* parameters. **(c)** The area under silhouette curve achieved by the consensus method using different values of *C* and *d*. The silhouette curve was generated by calculating the average silhouette value using the cluster R package for a set of clusterings obtained by varying the number of consensus clusters (*K*) from 2 to 50. The area under the silhouette curve was calculated using the DescTools R package.

The analysis was performed using $k$ = 15, and $r$ = 5 parameter values with the pbmc3k dataset. The data points depict average values across 200 ICP runs (*L = 200)*. The missing value at $C$ = 0.1, $d$ = 0.1 occurred due to the ICP algorithm becoming unstable when the number of clusters $k$ in some of the $L$ ICP runs started to decrease.

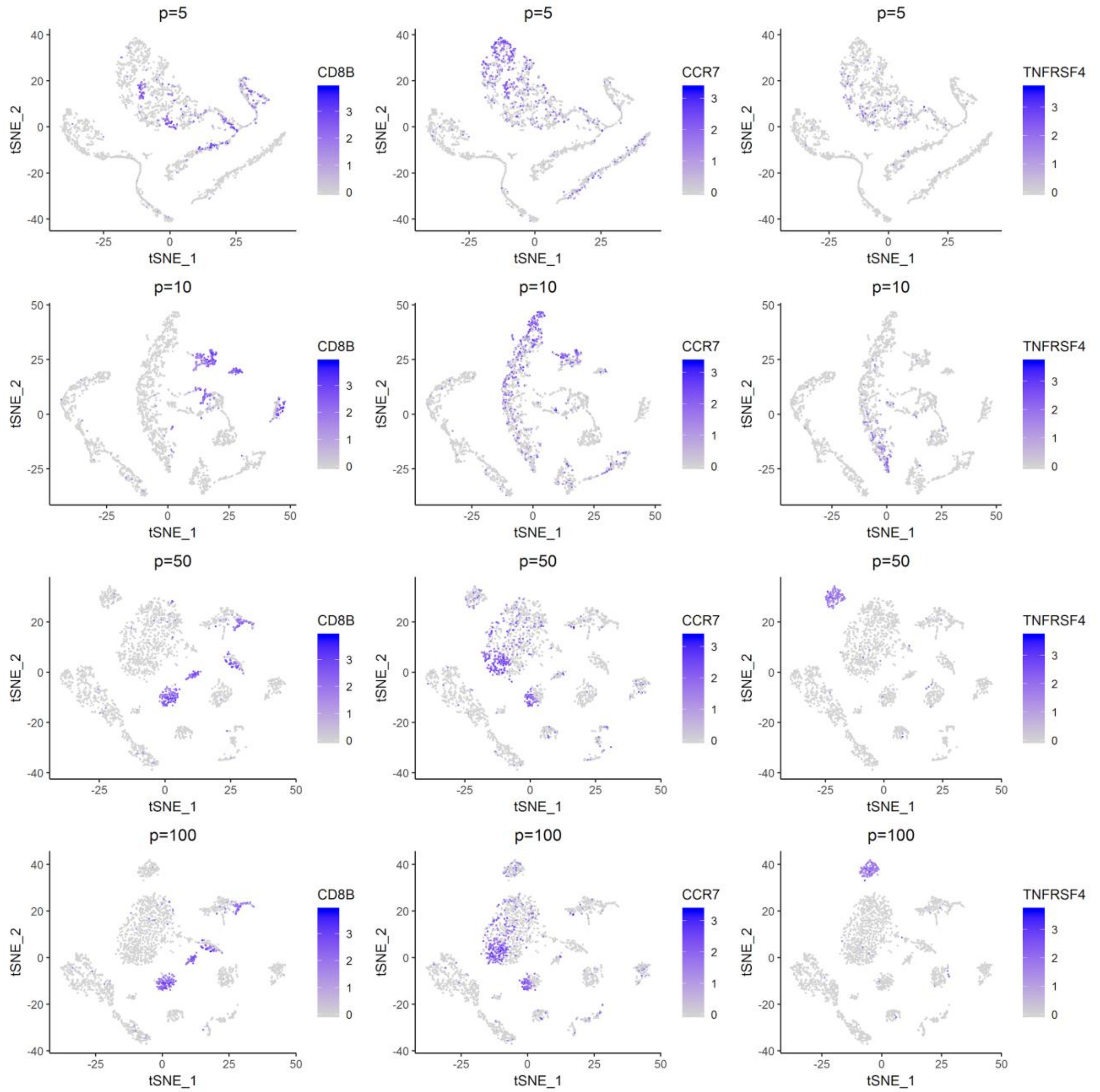


**Supplementary Figure 12**

Effect of the parameter *p* on the location of T cell subpopulations in t-SNE dimensionality reduction

*t*-distributed stochastic neighbor embedding (t-SNE) plots generated by ILoReg for the pbmc3k dataset with different values of the number of principal components (*p)* used in creating the consensus matrix from the joint probability matrix. The expression levels of three genes are highlighted: *CD8B* (CD8+ T cells), *CCR7* (naive T cells), *TNFRSF4* (activated CD4+ T cells). The other parameters were fixed: *d* = 0.3, *C* = 0.3, *k* = 15, *r* = 5, *L* = 200.

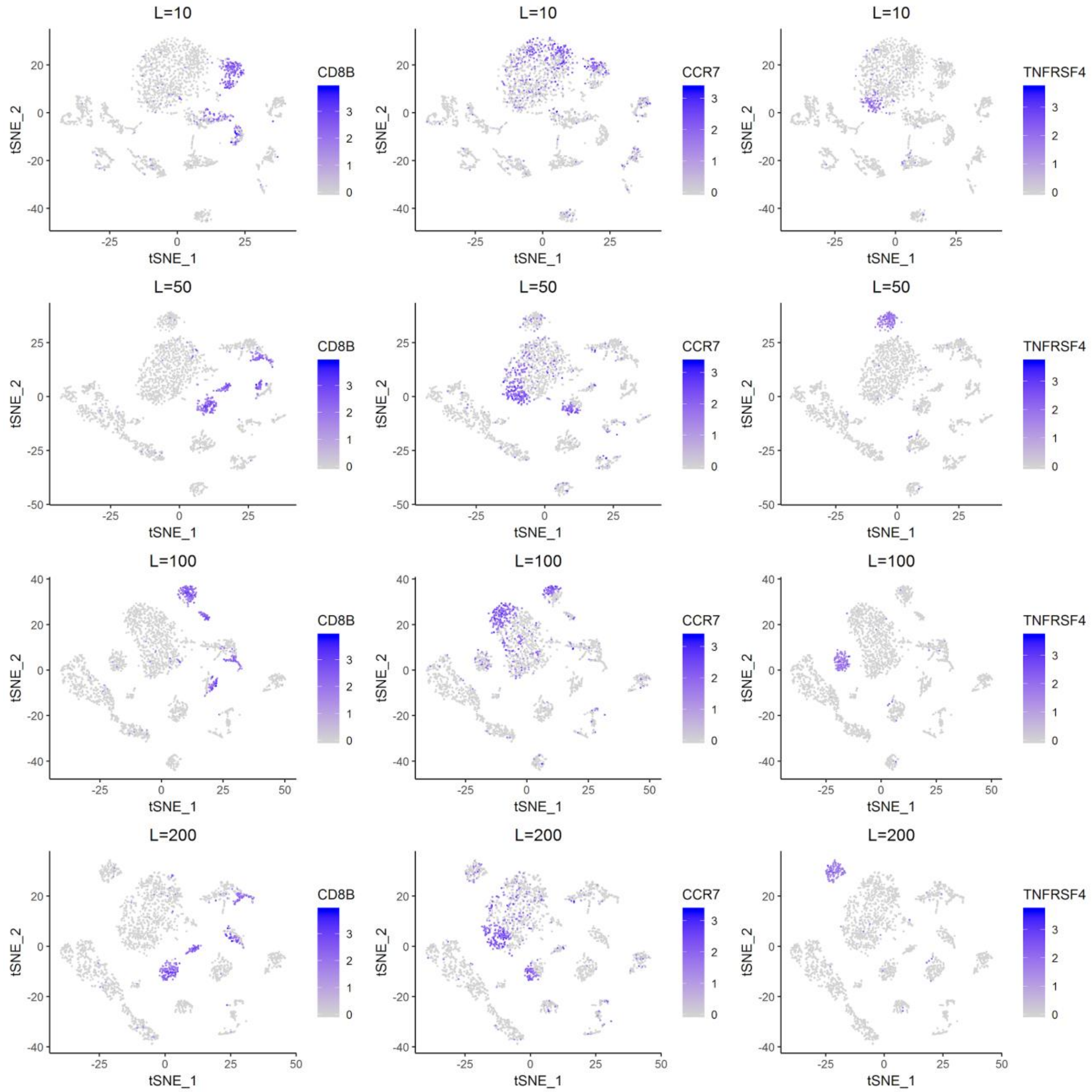


**Supplementary Figure 13**

Effect of the parameter *L* on the location of T cell subpopulations in t-SNE dimensionality reduction

*t*-distributed stochastic neighbor embedding (t-SNE) plots generated by ILoReg for the pbmc3k dataset with different values of the number of iterative clustering projection (ICP) runs (*L*). The expression levels of three genes are highlighted: *CD8B* (CD8+ T cells), *CCR7* (naive T cells), *TNFRSF4* (activated CD4+ T cells). The other parameters were fixed: $d$ = 0.3, $C$ = 0.3, $k$ = 15, $r$ = 5, $p$ = 50.

### Supplementary Results 1

#### Analysis of the pbmc3k dataset

We investigated in more detail peripheral blood mononuclear cell (PBMC) populations found by ILoReg from the pbmc3k dataset (**Fig. 2c**). In the t-SNE and UMAP representations that are visualized below, we have named cell populations found using the default parameter configuration of ILoReg ($k = 15$, $d = 0.3$, $C = 0.3$, $r = 5$, $L = 200$, $p = 50$).

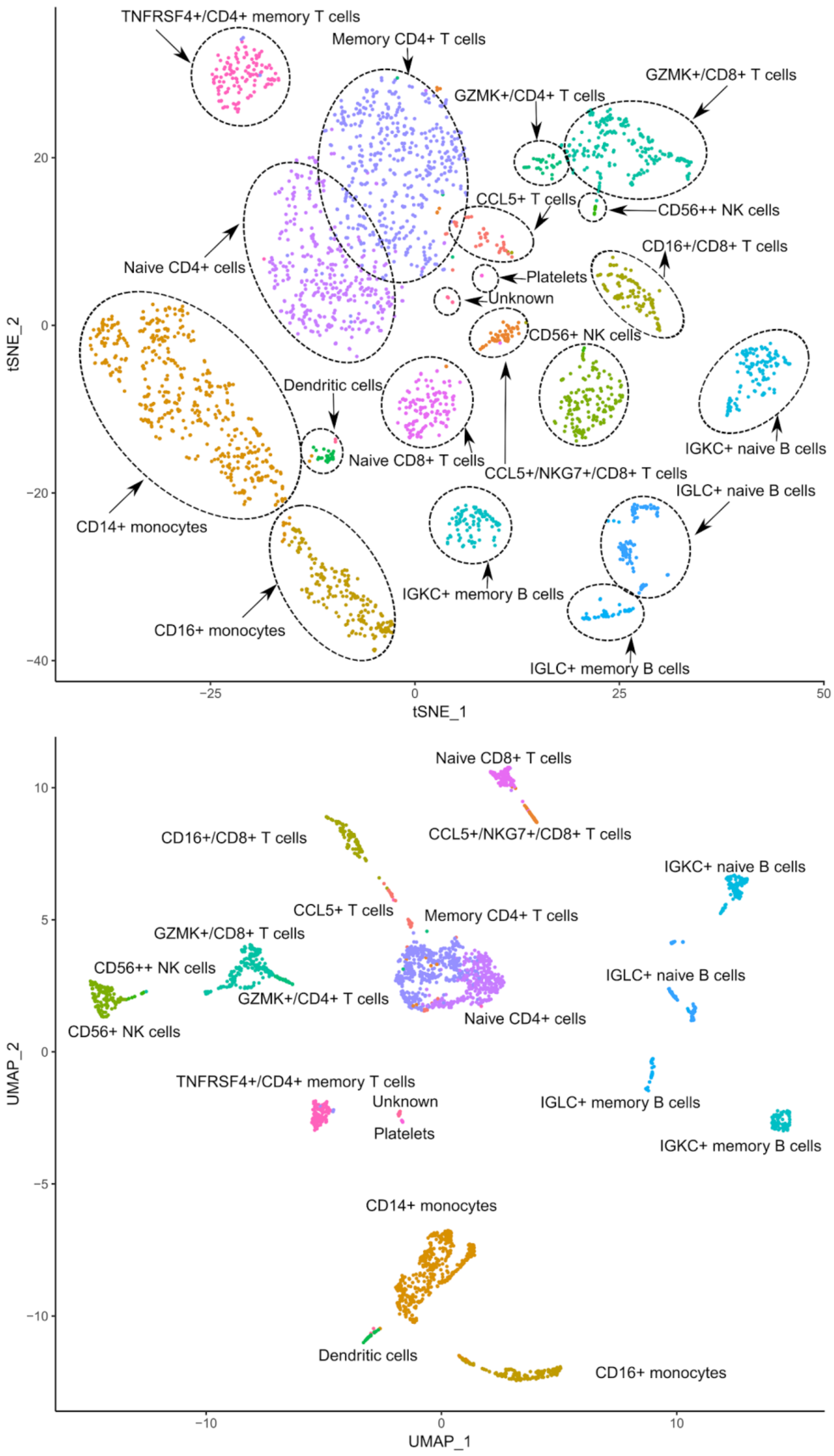


t-SNE and UMAP representations generated by ILoReg for the pbmc3k dataset showing the cell types named.

Expression of the two CD8 genes (*CD8A* and *CD8B*) and *CD4* are commonly used to differentiate CD8+ and CD4+ T cells, whereas *CCR7* and *S100A4* has been shown to be upregulated in naive and memory T cells, respectively. Therefore, the *CD3D*+/*CD8*-/*CCR7*+/*S100A4*- and *CD3D*+/*CD8*+/CCR7+/*S100A4*- cell populations correspond to naive CD4+ and CD8+ T cells[2,3], respectively. The second part of the large CD4+ T cell population contains *S100A4*+/CCR7-/*CD4*+ cells, bearing similarity to memory CD4+ T tells. To provide further support for the classification of naive and memory CD4+ T cells, the putative naive CD4+ T cell population had a higher expression of *FHIT,* a gene that has been shown to be upregulated in naive CD4+ T cells, whereas *TNFRSF4* and *AQP3* have been shown to be upregulated in T cells that have differentiated beyond the naive state[,29], such as memory T cells. Interestingly, the *TNFRSF4*+ cells formed a distinct disconnected cluster from the rest of the memory CD4+ T cells. In addition to the naive CD8+ T cells, four other CD8+ T cell populations are visible in the t-SNE plot. The *CCL5*+/*NKG7*+ population located adjacent to the naive CD8+ population did not express any granzymes (*GZMB*, *GZMA*, *GZMK*, *GZMH*), whereas the population close to the NK cells expressed *GZMB*, *GZMA* and *GZMH*, but also *FCGR3A*. *FCGR3A*+ CD8+ T cells have been previously characterized as an intermediate state in the developmental process from naive CD8+ T cells to *GZMK*+ CD8+ T cells[6]. A part of the *GZMK*-expressing population is depleted in expression of both *CD4* and the two CD8 genes, indicating it consists of double-negative T cells, such as MAIT or gamma delta T cells[7,8].

ILoReg identified two natural killer (NK) cell populations that are in agreement with the CD56 dim (CD56+) and CD56 bright (CD56++) NK cell types that have previously been identified using cellular indexing of transcriptomes and epitopes by sequencing (CITE-seq)[29]. In both experiments CD56++ cells expressed *GZMK*, but not *FCGR3A* (*CD16*).

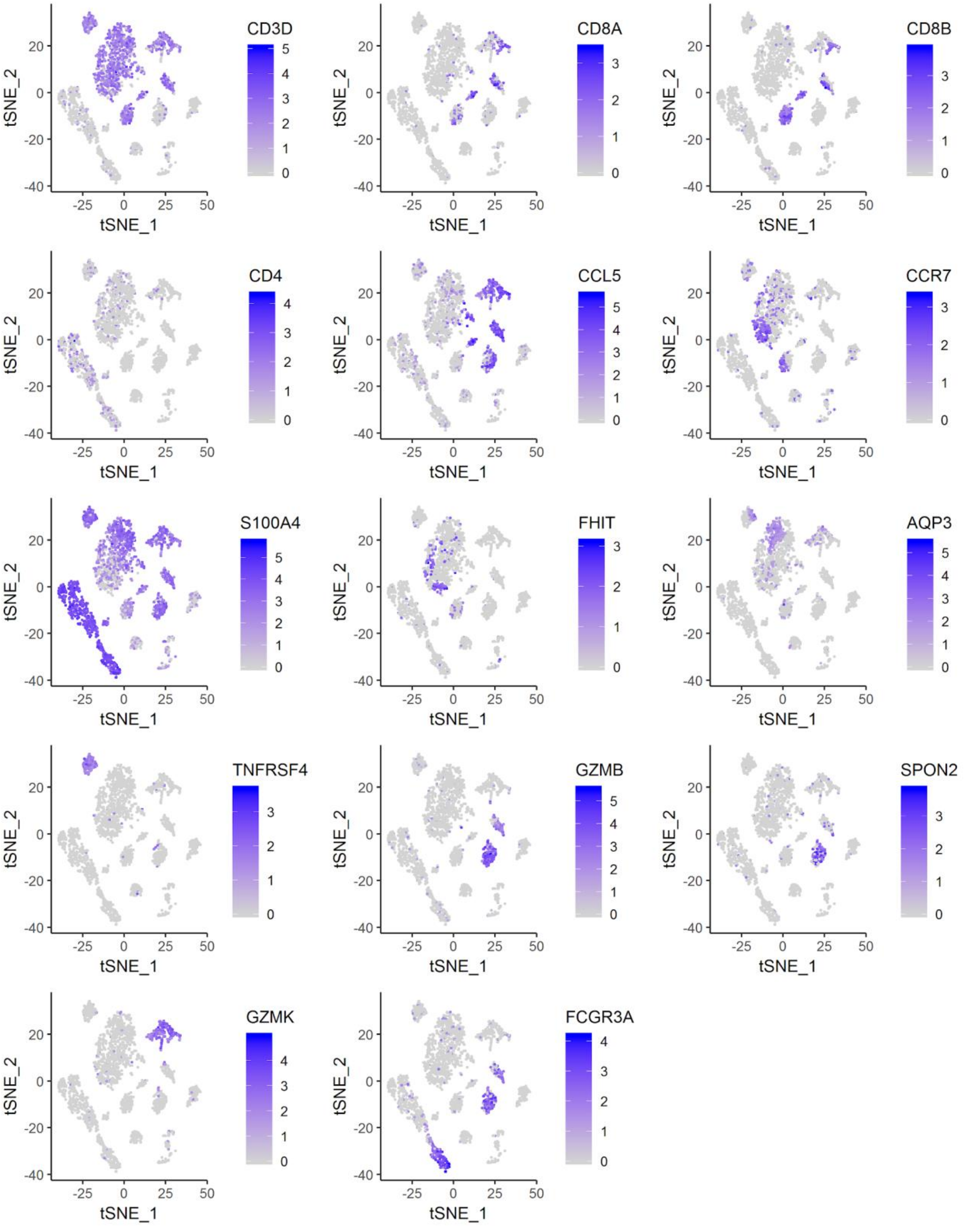


t-SNE plots generated by ILoreg highlighting expression levels of T and NK cell gene markers: *CD3D* (T cells), *CD8A* (CD8+ T cells), *CD8B* (CD8+ T cells), *CD4* (CD4+ T cells), *CCL5* (activated CD8+ T cells), *CCR7* (naive T cells), *S100A4* (effector and memory T cells), *FHIT* (naive CD4+ T cells), *AQP3* (CD4+ memory T cells), *TNFRSF4* (*TNFRSF4+* CD4+ T cells), *GMZB* (CD8+ effector T cells and CD56+

NK cells), *SPON2* (CD56+ NK cells), and *GZMK* (CD56++ NK cells, CD8+ effector T cells and double-negative T cells), *FCGR3A* (CD56+ NK cells and CD8+ effector T cells).

ILoReg partitioned the B cells into several distinct subpopulations. *TCL1A*+/*CD27*- and *TCL1A*-/*CD27*+ B cells correspond to naive and memory B cells[7,9], respectively. Further B cell populations of interest include populations with differentially expressing genes coding immunoglobulin light chain peptides (*IGLC2*, *IGLC3* and *IGKC*), being agreement with the classification of B cells into lambda (*IGLC2*+/*IGLC3*+) and kappa (*IGKC*) light chain carrying B cells[10].

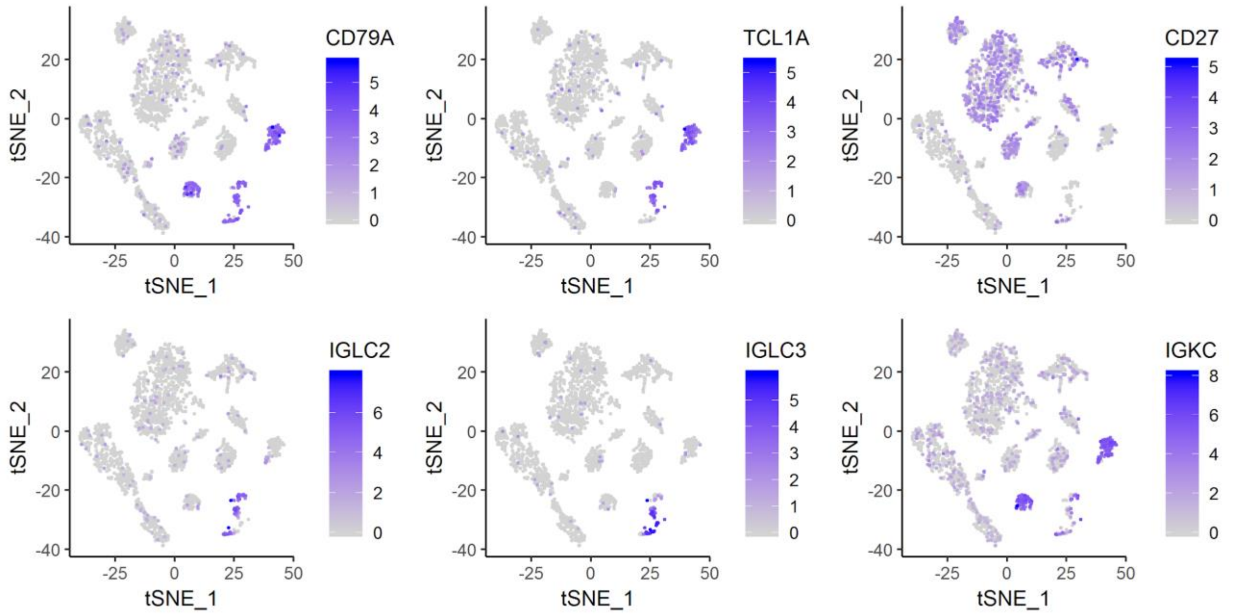


t-SNE plots generated by ILoreg highlighting expression levels of B cell gene markers: *CD79A* (all B cells), *TCL1A* (naive B cells), *CD27* (memory B cells), and *IGLC2*, *IGLC3* (B cells with lambda light chain), *IGKC* (B cells with kappa light chain).

ILoReg identified the two most common monocyte subtypes: CD14+ (classical) and CD16+ (non-classical) monocytes[11].

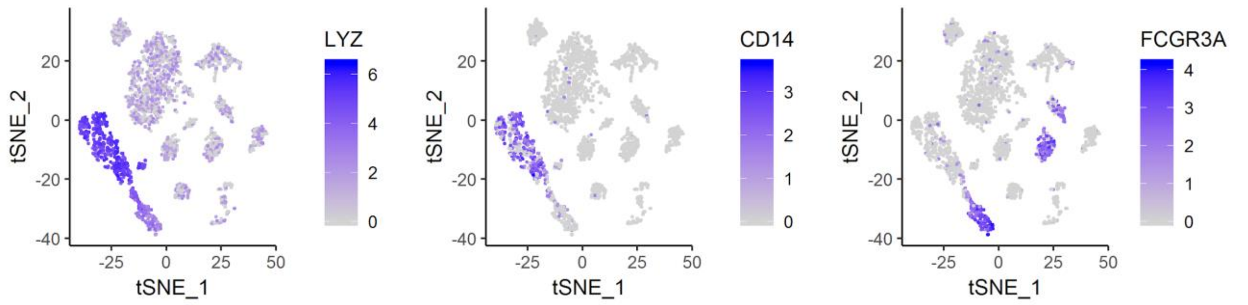


t-SNE plots generated by ILoreg highlighting expression levels of monocyte gene markers: *LYZ* (all monocytes), *CD14* (CD14+ monocytes), *FCGR3A* (CD16+ monocytes).

Finally, ILoReg identified rare PBMC cell types: dendritic cells (FCER1A+)[11] and platelets (GP9+/PPBP+)[12].

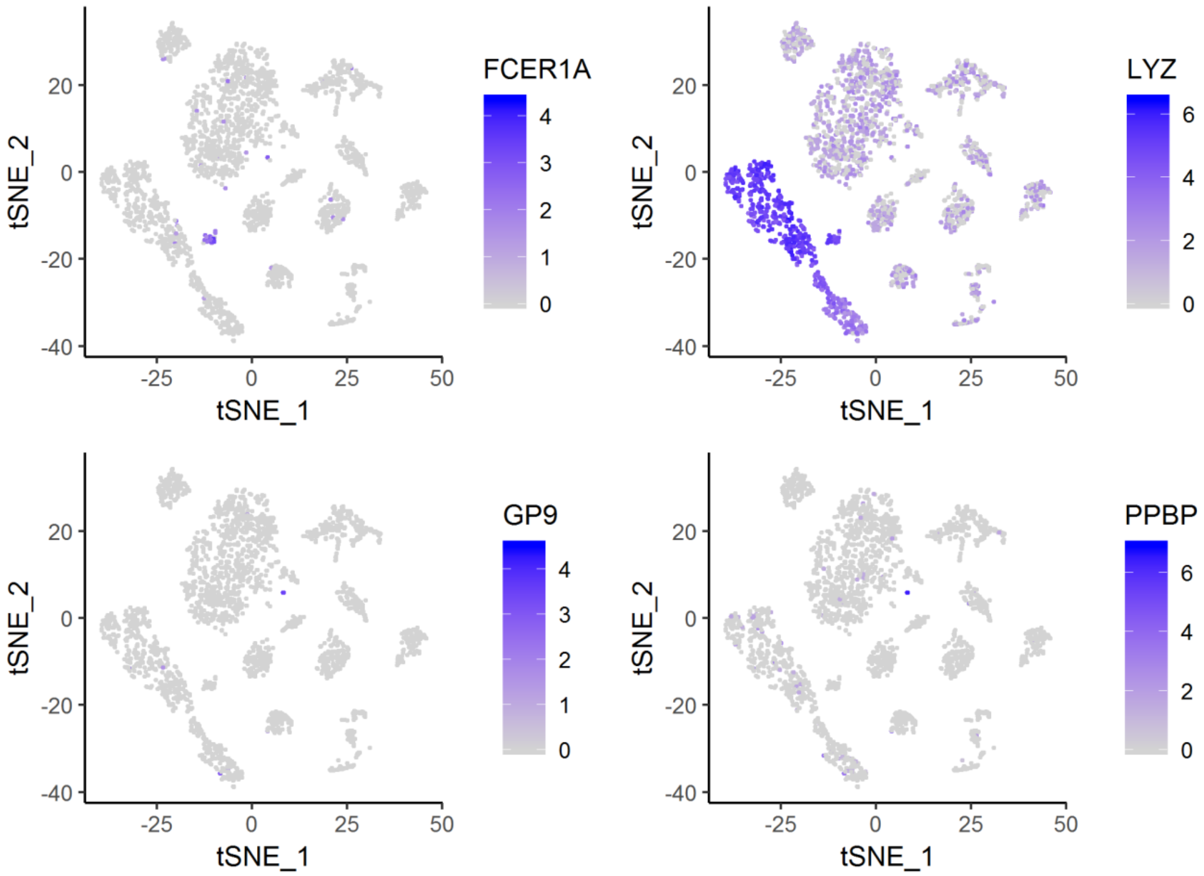


t-SNE plots generated by ILoReg highlighting expression levels of dendritic cell and platelet gene markers: *FCER1A* (dendritic cells), *LYZ* (monocytes and dendritic cells), *GP9* (platelets) and *PPBP* (platelets).

To identify gene markers for the clusters, we performed differential expression analysis (**Methods**). We visualized the top 10 gene markers for all the clusters selected based on log2 fold-change of the average expression values. The genes were filtered with 0.05 and 0.5 thresholds of the Bonferroni adjusted p-value and log2 fold-change, respectively. The full results of the comparison are in **Supplementary File 2**.

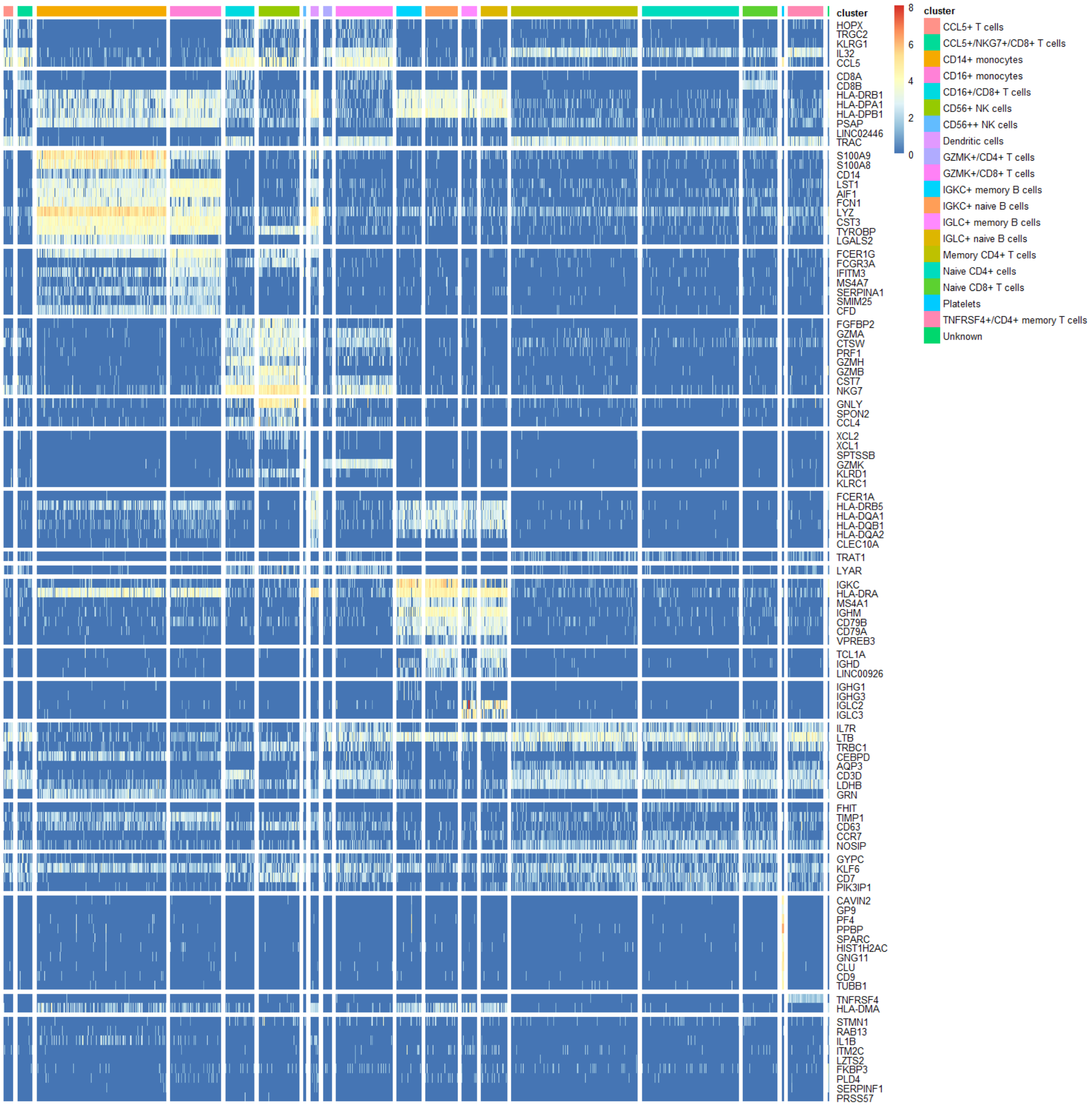


Gene marker heatmap of the top 10 genes that were selected based on log2 fold-change. To notify, for some clusters fewer than 10 statistically unique genes were found, and some genes were DE in multiple clusters.

### Supplementary Tables

**Supplementary Table 1**

Clustering algorithms used in benchmarking.

| Name | Clustering workflow | 2D visualization workflow | Method for estimating the optimal number of clusters (*k*) | Version |
|---|---|---|---|---|
| ILoReg | ICP *L* times + PCA + hierarchical | ICP *L* times + PCA + t-SNE or UMAP | Silhouette | 0.1.0 (Git reference ID “85196be6”) |
| Seurat[7] | Feature selection + PCA + graph-based | Feature selection + PCA + t-SNE, UMAP etc. | None (*k* by default resolution value 0.8) | 3.0.0 |
| SC3[8] | Feature selection + Distance matrices with three measures + PCA + k-means + CSPA + hierarchical | None (Feature selection by SC3 + PCA + t-SNE by Rtsne R package via scater R package) | Random matrix theory | 1.12.0 |
| CIDR[9] | Imputation + PCA + hierarchical | Feature selection + PCA | Calinski-Harabasz Index | 0.1.5 |
| RaceID3[10] | Feature selection + *k*-medoids | Feature selection + t-SNE or kNN graph | First *k* for which the decrease in within-cluster dispersion does not change (saturation) | 0.1.3 |

**Supplementary Table 2**

Summary of the datasets used in benchmarking.

| Study | Organism | Number of cells per dataset | Number of clusters per dataset | Protocol | Units | Standard |
|---|---|---|---|---|---|---|
| Pollen[13] | human | 301 | 11 | SMARTer | TPM | Gold |
| Baron[14] | human | 1937,1724, 3605,1303 | 14 | inDrop | UMI | Silver |
| Galen[15] | human | 108,188, 643,3738,1 431,807 | 13,15,15,1 5,9,5 | Seq-Well | UMI | Silver |
